# Evolution of cell-to-cell variability in stochastic, controlled, heteroplasmic mtDNA populations

**DOI:** 10.1101/072363

**Authors:** Iain G. Johnston, Nick S. Jones

## Abstract

Populations of physiologically vital mitochondrial DNA (mtDNA) molecules evolve in cells under control from the nucleus. The evolution of populations of mixed mtDNA types is complicated and poorly understood, and variability of these controlled admixtures plays a central role in the inheritance and onset of genetic disease. Here, we develop a mathematical theory describing the evolution and variability in these stochastic populations for any type of cellular control, showing that cell-to-cell variability in mtDNA, and mutant load, inevitably increases with time, according to rates which we derive and which are notably independent of the mechanistic details of feedback signalling. We show with a set of experimental case studies that this theory explains disparate quantitative results from classical and modern experimental and computational studies on heteroplasmy variance in different species. We demonstrate that our general model provides a host of specific insights, including a modification of the often-used but hard-to-interpret Wright formula to correspond directly to biological observables, the ability to quantify selective and mutational pressure in mtDNA populations, and the pronounced variability inevitably arising from the action of possible mtDNA quality-control mechanisms. Our general theoretical framework, supported by existing experimental results, thus helps understand and predict the evolution of stochastic mtDNA populations in cell biology.

## Introduction

Molecules of mitochondrial DNA (mtDNA) form dynamic evolutionary populations within cells, replicating and degrading according to cellular control signals [1, 2]. MtDNA can vary due to mutation or artificial manipulation [3]; the proportion of mutant mtDNA in a cell is referred to as heteroplasmy. MtDNA encodes vital aspects of the bioenergetic machinery of eukaroytic cells; mtDNA variability can thus have dramatic cellular consequences, including devastating genetic diseases and numerous other conditions [3], making a theoretical understanding of this complex evolutionary system important. Understanding the natural feedback control acting on mtDNA populations is also a vital step in the development of artificial approaches to control mitochondrial behaviour with genetic tools [4, 5]. The population variances of mtDNA types, and heteroplasmy variance, are of particular importance, owing to their implications for maternal transmission of dangerous mutations [6] and the manifestation of pathologies dependent on the range of heteroplasmies present in a tissue [7], including the demonstration that a very small proportion of cells exceeding a heteroplasmy threshold can lead to pathologies [8].

Stochasticity underlies cell biology; cellular processes including gene expression [9, 10, 11], DNA replication [12], and mitochondrial and mtDNA dynamics [13, 14, 15, 16] are subject to fundamentally stochastic influences. Variability in mitochondria can be a leading contributor to cell physiological behaviour, making mitochondria an important target for explanatory stochastic models [15]. Existing studies have included stochastic modelling and numerical treatments of mitochondrial [17] and mtDNA populations with the assumption of specific control mechanisms [18, 19, 1, 14, 20]. Other theoretical studies have drawn on classical statistical genetics, notably including the well-known Wright formula [21, 22], to produce a description of partitioning at cell divisions, but the role of stochastic mtDNA dynamics between cell divisions is largely omitted. Although recent experimental studies are starting to shed light on cellular control of mtDNA [14, 16], a general theoretical framework is currently absent. Here we address this open question by constructing a general, bottom-up stochastic description of mtDNA populations subject to arbitrary cellular control mechanisms, providing analytic results for the predicted behaviour associated with any mtDNA control mechanism, and adapting the classic Wright formula to account for and interpret stochastic mtDNA dynamics. Notably, our approach and results hold independently of the details of specific regulatory mechanisms underlying mtDNA feedback signalling, providing a general theoretical framework for control of stochastic mtDNA populations across different species and environments.

As we develop the theoretical framework to address mtDNA dynamics below, we will consider a set of applications of the theory, linking with existing experimental data from a variety of studies to validate our approach and obtain quantitative results and predictions on the processes governing mtDNA dynamics. We will focus on two questions arising from the study of mtDNA diseases: (a) how and at what rate does cell-to-cell heteroplasmy variance increase; and (b) how does selective pressure against a particular mtDNA mutation affect cellular mtDNA populations. The single-cell measurements required to address these questions directly remain challenging: we aim to show that mathematical theory, appropriately validated and refined with available data, allows us to make quantitative progress understanding this important behaviour. Fig. 1 summarises the central biological messages arising from development and analysis of our theory.

**Figure 1:**
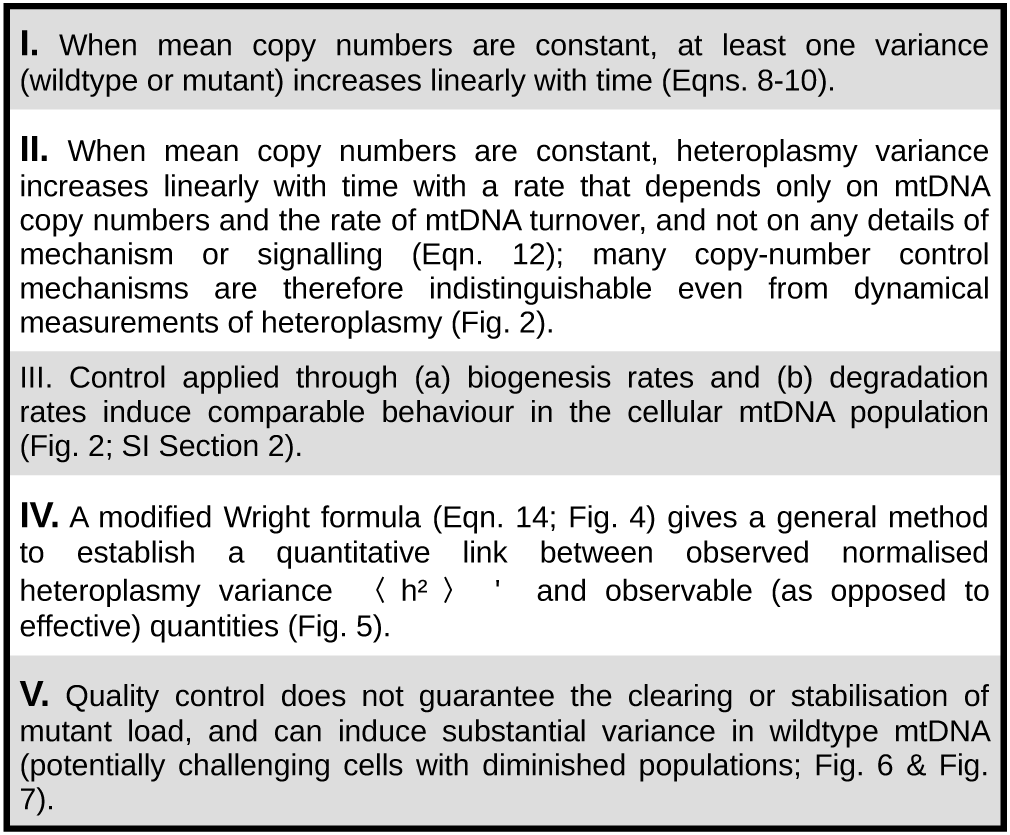
Key biological findings from our mathematical model. These results arise when the assumptions of our modelling approach hold – that cells are heteroplasmic, mtDNA replication and degradation are Poisson processes with the same rates for mutant and wildtype mtDNA, and feedback mechanisms depend only on the current state of the system. We justify these assumptions with reference to experimental data in the text and in Appendix A1.

## Methods

We will consider mtDNA populations in cells that are heteroplasmic with two non-recombining haplotypes, though this treatment can readily be extended to other species. We write a state with *w* wildtype mtDNAs and m mutant mtDNAs as {*w, m*}. We first consider the class of systems where both haplotypes are subject to the same degradation rate *ν* and the same replication rate *λ*, both of which may be general functions of both haplotype copy numbers. This model thus represents the situation where no direct selective difference exists between mutant and wildtype. This assumption only holds for some biological cases (see Ref. [23] and references therein for a review of studies where mtDNA types segregate unevenly) and will be relaxed later. We also assume that cellular control is based only on the current state of the cellular mtDNA population, and not its history. The dynamics governing the system then consist of a set of Poisson processes:

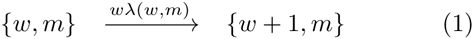

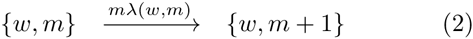

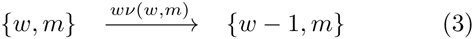

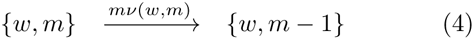

This formalism captures a wide range of models for mtDNA dynamics (see below). We will begin with the assumption that the system does not undergo cell divisions, and has a stationary state in the population mean of both haplotype copy numbers, and will write this steady state as 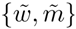. This initial picture is more appropriate for quiescent cell types or mtDNA ‘set points’ than for the pronounced changes in mtDNA copy number that occur during development [2, 14]. We will later generalise this picture to allow for arbitrary changes in copy number.

We will first consider general results from this formalism, applicable to a wide variety of possible cellular behaviours. We will then illustrate its application with a range of previously proposed, and new, feedback mechanisms.

## Results

### Copy number variance with stable population means

Any control mechanism of the form Eqns. 1-4 (including manifestations of feedback control) can be represented to linear order by a Taylor expansion of its rates about 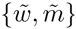 (the steady state exists by construction with our previous assumption):

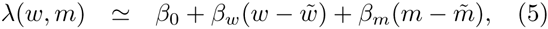

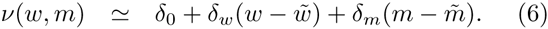

It will readily be seen that to support a stable population mean at 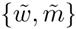, *δ*_0_ = *β*_0_. Assuming that *w* and *m* can be written as the sum of a deterministic and a fluctuating component, we use Van Kampen’s system size expansion to find a Fokker-Planck equation describing the behaviour of *w* and *m* governed by Eqns. 5-6 [24, 25]. From this equation we extract expressions for the time behaviour of the mean and variance of *w* and *m* (see SI Sections 1-2).

We show in SI Section 4 that attempting to identify a stable state for population variances and covariance yields the condition:

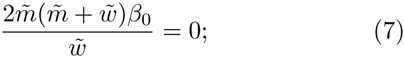

hence, any population mean state in which mutant content 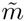 is nonzero does not admit a stationary solution for variances, unless *β*_0_ = 0. If *β*_0_ = 0 then there is no further change to the system once steady state has been reached (no stochastic turnover occurs) and the system remains frozen thereafter. In other words, *for a nonzero mutant population and nonzero mtDNA turnover, the variance of at least one mtDNA population will change with time.*

The Fokker-Planck equation can be used to compute the expected behaviours of 〈*w*^2^〉 (wildtype variance), 〈*m*^2^〉 (mutant variance), and 〈*wm*〉 (wildtypemutant covariance) for a given control mechanism. The variance and covariance solutions display some transient behaviour, involving terms on the timescale 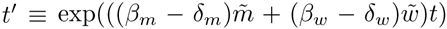. As, for stability, *β_i_* are nonpositive and *δ_i_* are nonnegative, *t*′ is either a constant or an exponentially decaying function of time *t*. The expressions thus subsequently converge to linear trends for large *t*:

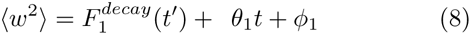

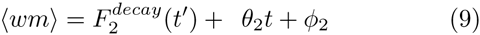

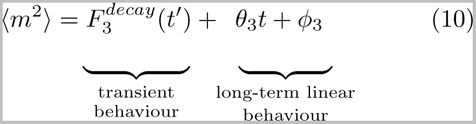

The forms of the transient functions 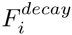, and the constants *θ*_*i*_ and *ϕ*_*i*_ are given in SI Section 2 and are functions only of the difference between replication and degradation rates (*β*_*i*_ − *δ*_*i*_), steady-state copy numbers 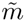 and 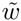, and mitophagy rate *β*_0_. Furthermore, the structure of these expressions is such that for *β*_0_ ≠ 0 and nonzero *w* and *m,* at most one of the *θ*_*i*_ can be zero, *θ*_1_ ≥ 0, and *θ*_3_ ≥ 0. Thus, around the mean 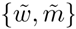, *either wildtype variance or mutant variance, or both, increase linearly with time (Fig. 1 I).* As time continues, the increasing variance means that extinction of one mtDNA becomes increasingly likely: implications of this behaviour are explored below.

The mathematical structure of the solutions only ever involves the *difference* between replication and degradation rates (*β*_*i*_ − *δ*_*i*_), showing that *control of (a) biogenesis rates and (b) degradation rates induce comparable behaviour in the cellular mtDNA population (Fig. 1 III)*.

### Heteroplasmy statistics

As shown in SI Section 5, a first order Taylor expansion of a function of random variables gives an approximation for the variance of *h* = *m*/(*w* + *m*):

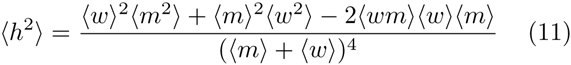

and we can then use previously obtained expressions for 〈*w*〉, 〈*m*〉, 〈*w*^2^〉, 〈*m*^2^〉, 〈*wm*〉 to compute this approximate heteroplasmy variance. Neglecting transient terms and using 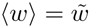 and 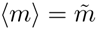 in Eqn. 11, gives, after some algebra,

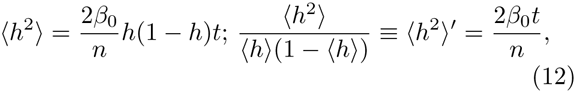

where 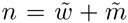, with 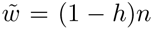 and 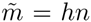: thus *h* is (mean) heteroplasmy and n is (mean) total copy number (recall that *β*_0_ = δ_0_ in steady state). In Eqn. 12 we have used normalised heteroplasmy variance 〈*h*^2^〉′, accounting for the dependence of 〈*h*^2^〉 on the magnitude of h; (h^2^)' is the quantity most often reported in experimental studies.

In other words, when the system size expansion is valid (see below), *for any control mechanism, heteroplasmy variance in the copy number steady state increases linearly with time with a rate that depends only on the copy numbers of the system and the timescale of random turnover (Fig. 1 II)* (Eqn. 12 is independent of the *β* and *6* terms in Eqns. 5-6). As we discuss later, this observation implies that many possible mechanisms could be responsible for the same observed trend in heteroplasmy variance, meaning that measurements of heteroplasmy variance alone, even if repeated at different time points, place only a limited mechanistic constraint on mtDNA dynamics [14]. In SI Section 7 we discuss experimental strategies that can more efficiently discriminate between different control mechanisms.

### Transient behaviour and cell divisions, and validity of the expansion

To obtain analytic insight, we have thus far focussed on modelling mtDNA behaviour using the system size expansion when a steady-state assumption had already been applied. Transient behaviour can also be explored by employing the system size expansion directly on the appropriate master equation, using the full expressions for λ(*w, m*) and *ν*(*w,m*) (see SI Sections 2 & 6). Relaxing the steady state assumption means that the ODEs describing variance behaviour are analytically intractable for many forms of λ(*w, m*), *ν*(*w,m*). However, they can simply be solved numerically and, as shown in subsequent sections, well match stochastic simulation (which of course is numerically far more intensive). This ODE approach fully accounts for non-equilibrium behaviour – including transient relaxation, cycling, and so on – while the system size expansion remains appropriate (see below).

This analysis can readily be used to characterise the effect of partitioning mtDNAs at cell divisions. To compute the time behaviour of variance where cell divisions occur at arbitrary times, we invoke a linear noise assumption [24, 25], first using the ODEs above to compute the variance behaviour within one cell cycle. Partitioning rules for copy number statistics are then applied, and the resulting post-partition statistics are used as the initial condition for a next phase of ODE solution. We here illustrate this process for binomial partitioning of mtDNAs to connect with recent studies in mice [14] and HeLa [15], although with an appropriate choice of partitioning rules, this approach can be used to address any partitioning regime (for example, the sub-binomial case recently reported in fission yeast [16]). In the case of binomial partitioning, the appropriate partitioning rules are 〈*w*〉 → 〈*w*〉/2, 〈*m*〉 → 〈*m*〉/2, 〈*w^2^*〉 → 〈*w*^2^〉/4 + 〈*w*〉/4, 〈*wm*〉 → 〈*wm*〉/4, 〈*m*^2^〉 → 〈*m*^2^〉/4 + 〈*m*〉/4, following straightforwardly from the variance of a binomial distribution with *p* = 1/2 as *n*/4. We will see below that this picture well describes the behaviour of stochastic populations in dividing cells: hence, *the total variance contributions of turnover between divisions and partitioning at divisions can be modelled as a linear sum*, and the behaviour of mechanisms across cell cycles is comparable to that within a cell cycle.

The results above hold for a nonzero mutant population. As copy number variance increases, we expect extinction of one mtDNA type to become increasingly likely. To address this behaviour, we must consider when the system size expansion itself, which is reliant on the validity of the linear noise approximation, holds. An important threat to this validity is a non-negligible extinction probability for one mtDNA type, whereupon a normal distribution no longer adequately models the copy number distribution. Heuristically, this situation arises when, for example, 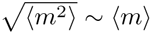. Another challenge arises due to the fact that, when *λ* and *ν* are functions of *w* and *m*, our linear theory is an approximation to the nonlinear dynamics that result. Highly nonlinear behaviour (for example, pronounced discrete steps in rates occuring at critical copy numbers) will therefore not be perfectly captured; but the ability of our theory to reproduce simulation of the fully nonlinear dynamics (in Figs. 2, 4, 6 and SI Section 3) suggests that the linear theory provides valuable insight into a wide range of biologically plausible behaviours. Treatments of fully nonlinear cases represent a substantial technical challenge which will be addressed in future work.

In these cases, the mean and variance of mtDNA populations are likely to be underestimated by the preceding analysis (see SI Section 3), and the heteroplasmy variance is overestimated, with the increase in 〈*h*^2^〉 with time gradually becoming sublinear. Fixation is also neglected by the deterministic version of the mean equations of motion, which allow an asymptotic descent to zero. Thus, the more general statement of our finding is that (i) for the period when extinction of either type is unlikely, variances and covariances change linearly (after transients); (ii) as extinction of one type becomes more likely due to this increased variance, the increasing trend continues but departs from those linear forms (in particular, the increase of 〈*h*^2^〉 slows to become sublinear); (iii) when extinction of one type is almost certain, the system tends towards its behaviour if only one type was present (ultimately stalling variance increase, as if 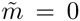 in Eqn. 7). The results we focus on in this main text can be viewed as describing the ‘quasistationary state’ where extinction is negligible; further quantitative details can be derived using, for example, adaptations of the system size expansion that address extinction [26].

### Specific control mechanisms and comparison with simulation

The previous results make no assumptions about the specific form of control applied to the mtDNA population, other than it depends only on current state and is manifest through the rates of Poissonian replication and degradation which are equal for both mtDNA species (and can be described with the system size expansion, as discussed above). We can exploit the generality of the preceding formalism to obtain results for any given (feedback) control mechanism, defined by a specific form of *λ*(*w, m*) and *ν*(*w,m*) in Eqns. 1-4.

We first consider the well-known ‘relaxed replication’ model [19, 1], which involves stochastic mtDNA degradation, coupled with mtDNA replication which is physically modelled as a deterministic process. We propose that, if degradation is regarded as a stochastic process (due to its microscopic reliance on complicated processes and colocalisations in the cell), picturing replication (which also relies on complicated interactions on the microscopic scale) as a stochastic process leads to a consistent stochastic generalisation (see Appendix A1). The corresponding model has exactly the same expressions for rates as in Ref. [1] (A in Fig. 2 i), but replication rate is now interpreted as the rate of a stochastic, rather than a deterministic, process.

**Figure 2:**
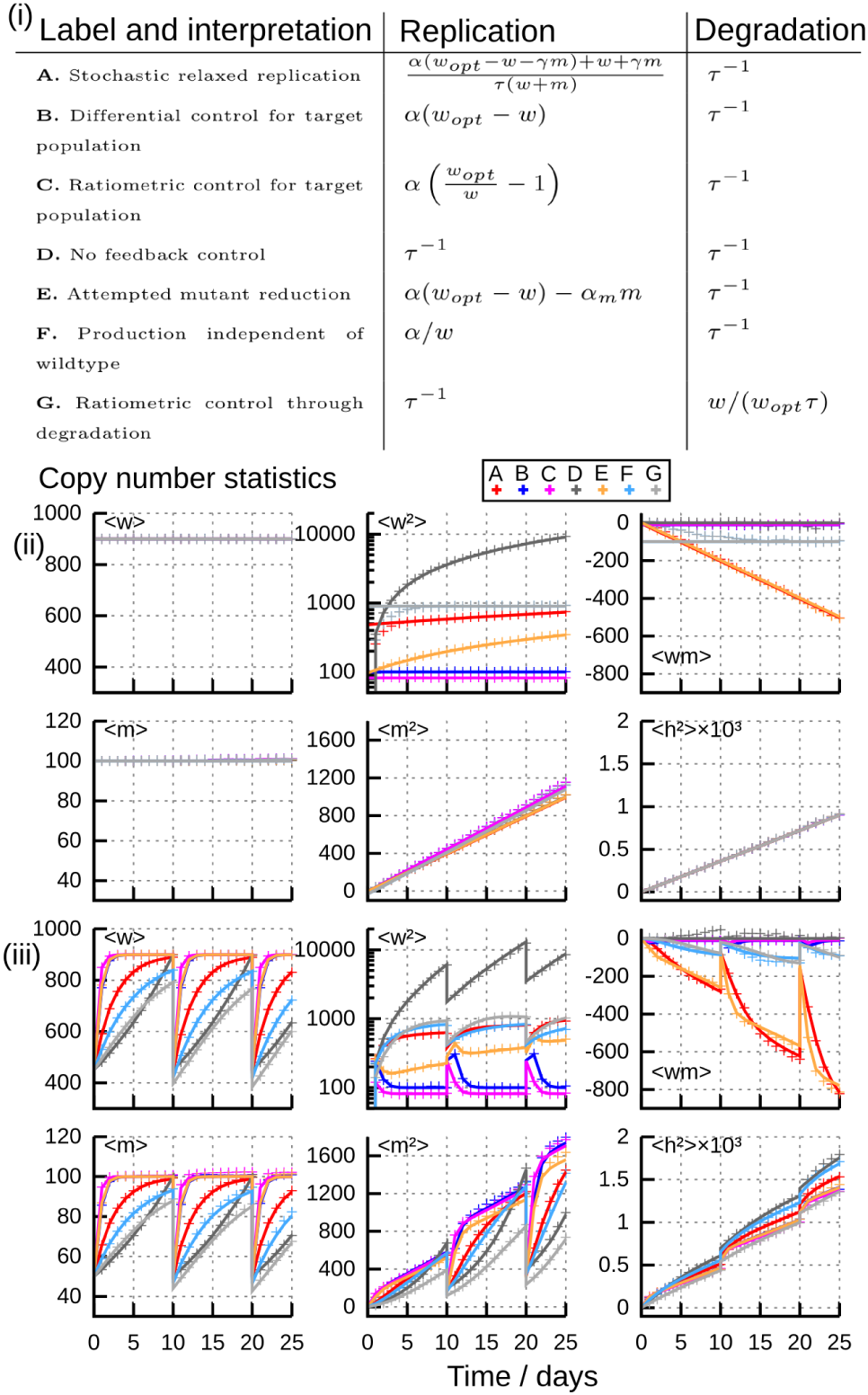
Behaviour of different specific control mechanisms. All control mechanisms have at least one population with time-increasing variance and so have increasing heteroplasmy variance; mean heteroplasmy 〈*h*〉 remains constant through these simulations. (i) Replication λ(*w,m*) and degradation *ν*(*w, m*) rates for model control mechanisms explored in the text, from existing studies and newly proposed here. (ii)-(iii) Copy number and heteroplasmy moments with time for these control mechanisms, (ii) in the absence of cell divisions and (iii) with binomial cell divisions every 10 time units. Analytic results (lines; full expressions in SI Section 3) match stochastic simulation (crosses) throughout. Parameters used are chosen to support the same steady state (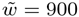, 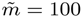) and with a turnover timescale of *τ* = 5 days. Other parameters: *γ* = 0, **α**_*m*_ = 0.001, **α** = 2 (except for models B, E, F, for which **α** = 0.002, 0.002, 200 respectively); 10^5^ stochastic simulations used.

We also introduce several other models for mtDNA control, to consider a range of potential functional forms, including differential and ratiometric control based on a target wildtype copy number, an absence of any feedback control, and others (Fig. 2 i B-G). The presence or absence of *w* and *m* in these expressions reflects what quantity is being sensed by the cell (wildtype mtDNA alone, mutant mtDNA alone, or a combination of the two). We further note that this general formalism can also incorporate physical constraints on the mtDNA population. For example, the hypothesis that mitochondrial *concentration* is controlled between cell divisions [15] (recently confirmed in fission yeast [16]) could correspond to *w_opt_*, the ‘target’ mtDNA number, being a linear function of cell volume in the models above; or could arise through passive birth-death dynamics (model D) with control implemented at the cell division stage (see previous section). The interpretations of these control mechanisms in terms of cellular sensing and the language of stochastic population processes are given in SI Section 3.

Fig. 2 ii illustrates the application of our analysis to these example control mechanisms in the absence of cell divisions. The close agreement between stochastic simulation and analytic results in steady state shows the generality of our theory. The long-term linear increases in one or both mtDNA variances are clear (Fig. 1 I), and trajectories of 〈*h*^2^〉 with the same steady state and turnover timescale are identical (Fig. 1 II). Fig. 2 iii, including cell divisions, demonstrates close agreement between ODE solutions and stochastic simulation, further showing that the linear noise treatment successfully captures stochastic behaviour over cell divisions. The close similarity of 〈*h*^2^〉 trajectories across divisions is a consequence of their aforementioned identity in steady state conditions with no cell divisions (Fig. 2 ii); differences are due to the difference in mechanism behaviour away from the steady state.

### Applications I: Heteroplasmy variance increases at constant mean copy number

The increase of heteroplasmy variance 〈*h*^2^〉 with time is of profound importance in determining the inheritance and onset of mtDNA diseases. Because disease symptoms often manifest only when heteroplasmy exceeds a certain threshold [7], increasing heteroplasmy variance with time can lead to pathologies even if mean heteroplasmy does not change (because a higher cell-to-cell variance implies a greater probability of a given cell exceeding a threshold) [14].

We sought experimental evidence to support the linear increase of 〈*h*^2^〉 predicted by our theory. Time course measurements of single-cell heteroplasmy values remain limited; we identified results from the *Drosophila* germline [27] and in the mouse germline for the NZB/BALB model [28, 29] and the HB model [14]. For these data, we compared the ability to fit the data of a null model (𝓗_0_ : 〈*h*^2^〉′ = **α** + ∊, where **α** is a constant), and an alternative model (𝓗_1_ : 〈*h*^2^〉′ = **α**+*βt*+∊), where 〈*h*^2^〉′ changes linearly with time as our theory predicts (Eqn. 12; Fig. 1 II). Using the Akaike information criterion (AIC) and assuming normally-distributed noise on mean 〈*h*^2^〉′ (ε ~ 𝒩(0, *σ*^2^), an assumption consistent with our linear approximation, but which can be further refined as in Ref. [29]), we found that the alternative, time-varying model was favoured in all cases, providing support for our theory (Fig. 3A-C).

**Figure 3:**
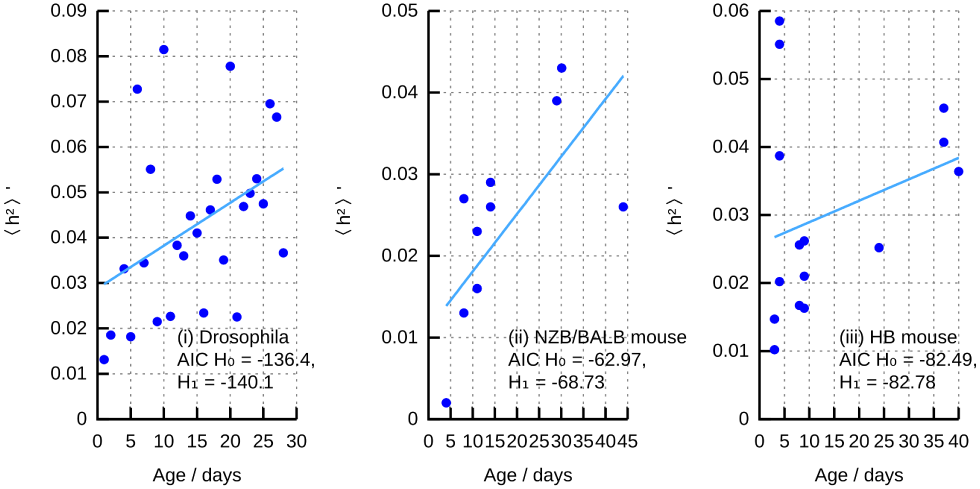
Experimental support for linear 〈*h*^2^〉 in crease. Normalised heteroplasmy variance 〈*h*^2^〉′ in model organism germlines over time (points); lines show maximum-likelihood linear fit for 〈*h*^2^〉 as a function of time. Insets give Akaike information criterion (AIC) values for 𝓗_0_(〈*h*^2^〉′ is constant) and 𝓗_1_ ((*h*^2^)′ increases linearly with time). Model organism and reference(s)): (i) *Drosophila* [27]. (ii) NZB/BALB mice [28, 29]. (iii) HB mice [14].

These results are quantitatively consistent with a previous study on the dynamics of heteroplasmy variance during the mtDNA bottleneck in mice [14] where a mechanism involving random mtDNA turnover and random mtDNA partitioning at cell divisions was found to best explain experimental observations. MtDNA in mice and rats often has a half-life of 10-100 days [23]. This corresponds to *β*_0_ = *δ_0_* = log 2/*t*_1/2_ = 0.03 – 0.003 days^−1^ and *τ* = *t*_1/2_/log 2 = 5 − 50 days. In Fig. 2 we show the patterns of copy number and heteroplasmy means and variances for *τ* = 5 days and 10^3^ mtDNA molecules per cell under different specific control strategies. The increase of 〈*h*^2^〉 from 0 to 10^−3^, corresponding for h = 0.1 to an increase in 〈*h*^2^〉′ from 0 to 0.011, matches the scale of change observed in the mtDNA bottleneck (though the bottleneck is complicated by changing population size n and compensatory changing turnover *β*_*0*_) [14]. Previous work has shown that the case with no feedback (D in Fig. 2) describes well the behaviour of mtDNA with cell divisions in mouse development [14]. In Appendix A1 we discuss further connections with previous theoretical studies; experimental cell-to-cell measurements in more quiescent cell types, while currently lacking, will provide valuable further tests of our theory.

### MtDNA turnover in the Wright formula

Powerful existing analyses of mtDNA population variance [22, 29] with widespread influence [3] have drawn upon a classical theory by Wright (and Kimura) [21, 30], describing stochastic sampling of a population of elements between generations. The resulting expression for expected heteroplasmy variance is the well-known equation sometimes referred to as the ‘Wright formula’ (though other equations also bear this name) [22, 3]:

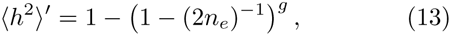

where *n_e_* is an effective population size and *g* is a number of generations. The mapping of this effective theory to the complicated mtDNA system is valuable to develop intuition but cannot capture the detailed dynamics of individual mtDNA molecules, due to assumptions (see Appendix A2) that mean the effective parameters of the theory (*n_e_* and *g*) cannot generally be interpreted as biological observables [22, 31], preventing quantitative analyses of mechanisms and dynamics [3]. In particular, *n_e_* does not generally correspond to a minimum mtDNA copy number (see Appendix A2), and, being a genetic rather than a physical parameter, ‘is unlikely ever to correspond closely to the number of anything’ [31].

The Wright formula, however, does accurately describe the heteroplasmy variance due to binomial sampling of 2*n_e_* real elements at cell divisions for an observable population size, and, as seen in the previous section, the additional effect of mtDNA turnover based on observable values can be included as an extra linear contribution. In general, this term will depend on the dynamics controlling the mtDNA population and can easily be calculated using the ODE approach above (Fig. 2).

In the case where no systematic change in mtDNA population size occurs with time, we can use the observation that 〈*h*^2^〉 trajectories are often comparable across a variety of different possible cellular control mechanisms (and identical in the steady state; Fig. 2 and Eqn. 12) to produce a simple approximate description linking 〈*h*^2^〉′ to observables. The simple steady state behaviour is given by Eqn. 12. To construct an approximation we use a simple estimate of mean population size over a cell cycle, writing 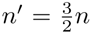, where *n* is the mtDNA population size immediately after division, and *n*′ thus gives a population size ‘average’ over the changes within a cell cycle. Using the above analysis with *w*_0_ = (1 − 〈*h*〉)*n*′, *m*_0_ = 〈*h*〉*n*′ (representing the ‘average’ populations of wildtype and mutant mtDNA), and *β*_0_ = 1/*τ* (so that *τ* is the timescale of mtDNA degradation), the corresponding expression in terms of *h* and *n* is then given by a ‘turnover-adjusted’ Wright formula (see Appendix A2):

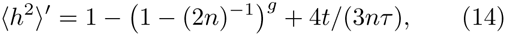

where *g* is the number of cell divisions that have occurred and t is the amount of time that has expired since an initial state with 〈*h*^2^〉′ = 0. This expression is subject to the conditions for system size expansion validity described above; thus, as fixation probability increases, the increase of 〈*h*^2^〉′ will drop below this prediction.

Fig. 4 illustrates the agreement between Eqn. 14 and stochastic simulation for the range of control mechanisms we consider under different population sizes and heteroplasmies. It is worth reiterating that more exact solutions for a given control mechanism can easily be computed using the preceding ODE approach, and stochastic analysis can also be used to quantitatively describe the effects of more specific circumstances (for example, the systematically varying population size through the mtDNA bottleneck [14]). In the case of no such systematic variation, and, crucially, *if the Poissonian model Eqns. 1-4 holds,* then Eqn. 14, *a modified Wright formula, represents a simpler, approximate way to establish a quantitative link between observed normalised heteroplasmy variance* 〈*h*^2^〉′ *and observable quantities (Fig. 1 IV)* – *n* (mtDNA copy number immediately after division), *g* (number of cell divisions), and *τ* (timescale of mtDNA turnover).

**Figure 4:**
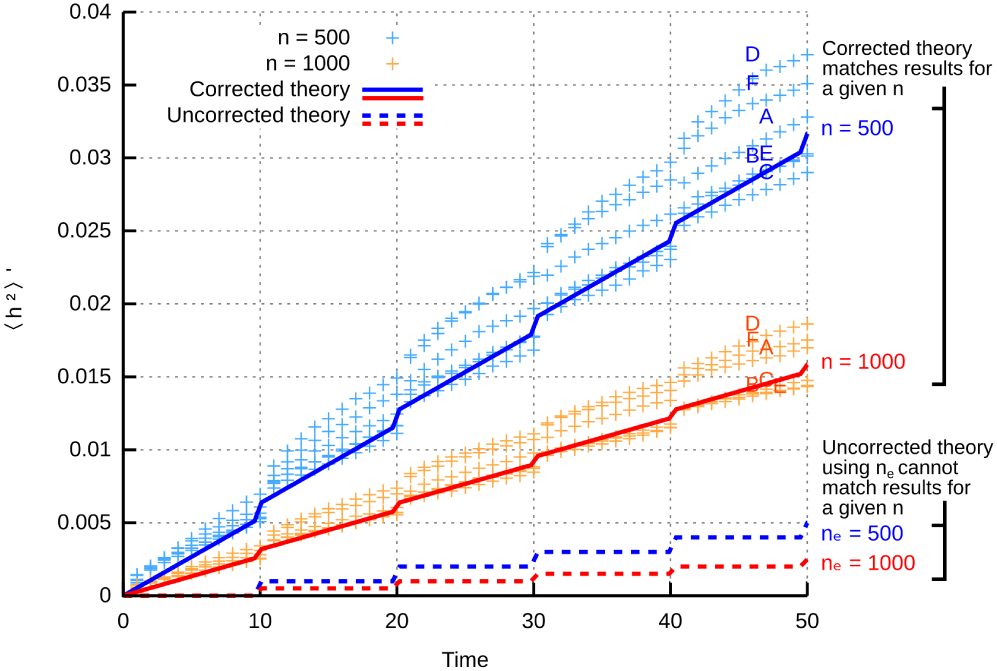
Correcting the Wright formula to include mtDNA turnover and connect to biological observables. Application of the Wright formula Eqn. 13 to stochastic mtDNA populations subject to division is only semi-quantitative: the Unadjusted Theory lines (dashed) for given effective population sizes *n_e_* do not describe the behaviour of simulations of mtDNA populations of that size (crosses). Correcting for mtDNA turnover with Eqn. 14 quantitatively connects Theory lines (solid) and simulation (crosses); further improvement can be achieved using the ODE approach in Fig. 2.

### Applications II: Linking physical and genetic rates with the modified Wright formula

The Wright formula is traditionally used to compare a heuristic, effective ‘bottleneck size’ across experimental systems (for example, in studies of different organisms [22] and of human disease [35]). In its uncorrected form this ‘bottleneck size’ can only be semi-quantitatively treated – bottleneck sizes can be ranked, but absolute values and differences cannot be straightforwardly interpreted. Our adaptation allows us to use this formula to connect the rates of physical subcellular processes with the resulting rates of genetic change.

To illustrate this connection, we focus on a particular period during mouse development. Between 8.5 and 13.5 days post conception (dpc) in the developing mouse germ line, cell divisions occur with a period of about 16 hours [36], giving g = 7 or 8 cell divisions in this period (of length *t* = 5 days). Copy number measurements during this period show that the mean total number of mtDNA molecules per cell remains of the order of n = 2000 (Fig. 5(i)) [28, 33, 34]. During this period, heteroplasmy variance 〈*h*^2^〉′ increases on average (but with substantial variability) from around 0.01 to 0.02 (Fig. 5(ii)) [32, 28]. Fig. 5(ii) shows a best-fit line to 〈*h*^2^〉′ data, with slope 1.52 × 10^−3^ day^−1^ (5-95% confidence intervals (1.11 – 1.92) × 10^−3^ day^−1^).

**Figure 5:**
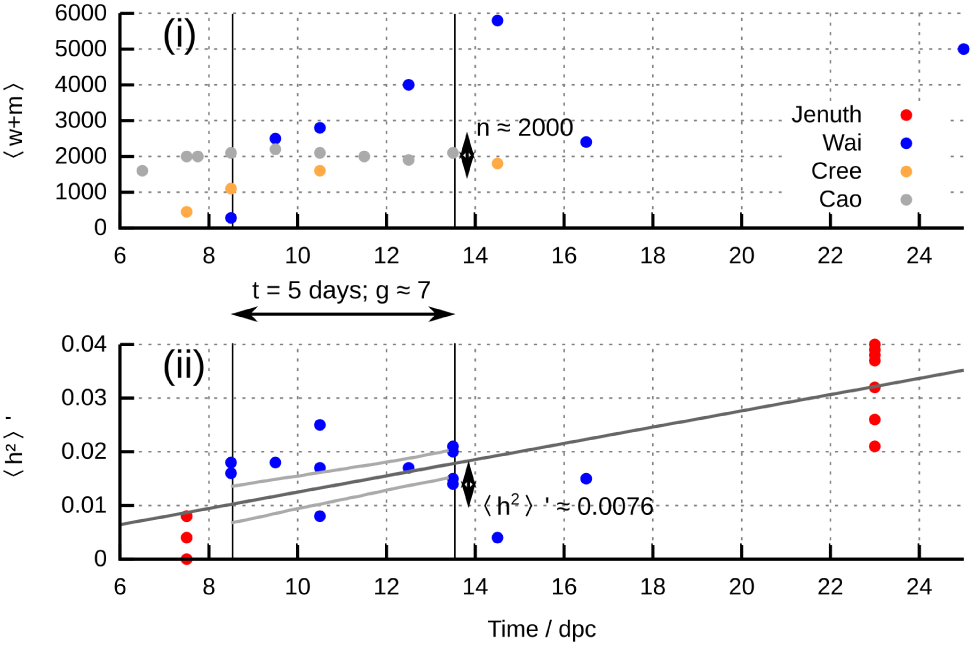
Using the adapted Wright formula to estimate mtDNA turnover. (i) MtDNA copy number n, taken to correspond to 〈*w* + *m*〉, in the mouse germ line during development. Data from several existing studies, referred to by first author [32, 28, 33, 34]. (ii) Normalised heteroplasmy variance 〈*h*^2^〉′ in this time period (points). Dark grey line is the mean inferred increase (which is linear, in agreement with our theoretical predictions, as in Fig. 3). Light grey lines give the 95% confidence intervals in the 8.5-13.5 dpc window, from a linear model fit. Values used in the turnoved-adjusted Wright formula (Eqn. 14) are given.

We can use these measurements in conjunction with the turnover-adjusted Wright fomula (Eqn. 14) to obtain estimates for the rate of mtDNA turnover during this period. Using Eqn. 14 with the best-fit 〈*h*^2^〉′ = 1.52 × 10^−3^ × 5 = 7.6 × 10^−3^, and *g* = 7 divisions, *n* = 2000 mtDNA molecules, *t* = 5 days gives the resulting estimate *τ* ≃ 0.57 days (5-95% confidence intervals 0.43-0.88 days, using the same values for *n, g, t*) for the characteristic timescale of mtDNA degradation. This increase in mtDNA turnover (relative to the *τ* ≃ 5 – 50 day timescale in differentiated tissues [23]) in the germline during this developmental period matches quantitative results from a more detailed study of the bottleneck reporting *τ* within the range 0.38-2.1 days (based on posteriors for *ν* = 1/*τ* between 0.02-0.11 hr^−1^) [14], and illustrates how a suitable mathematical model can be used to estimate biological quantities that are challenging to directly address with experiment [37].

### Influence of mutations, replication errors, and selective differences: quality control of replication errors

Our approach is easily generalised to include other processes than those described by Eqn. 1-4: in SI Section 6 we demonstrate that adding and changing appropriate processes allows us to analyse the effects on mtDNA mean and variance due to *de novo* mutations, replication errors, and multiple selective pressures. Our approach can be thus used to *characterise variability arising from selection and mutation under any control mechanisms, without requiring stochastic simulation.*

We can use this ability to explore a particular scientific question: if mtDNA replication errors occur and the cell attempts to clear the resulting mutant mtDNA through selective quality control, how do cellular mtDNA populations change? To investigate this question we introduce the process 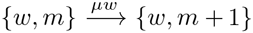 (replication errors – leading to the production of a new mutant mtDNA – occuring with rate μ) and reparameterise Eqn. 4 as 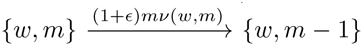 (a proportional increase of ∊ in mutant degradation compared to wildtype degradation). We thus model the situation where replication errors arise and the cell attempts to clear them through quality control, *while a control strategy for mtDNA populations is also in place.*

Fig. 6 illustrates the mean and variance of *w* and *m* in two different cases, distinguished by the relative magnitude of the selective difference (*∊*) and error rate (*μ*). This ratio is crucial in determining whether mutant mtDNA is cleared or increased: Fig. 6 (i) shows that mutant is cleared when (1 + *∊ν*) ≫ *μ* (selection is sufficiently strong to overcome errors), but when selective difference ∊ is insufficiently high, mutant mtDNA mean and variance (and heteroplasmy) increase with time. In both cases, we also observe substantial differences in mtDNA behaviour depending on the control model in place. Control models lacking an explicit target copy number (D (no feedback); F,G (immigration-like)) experience substantial increases in wildtype variance while mutant is being removed. Models involving a target wildtype copy number and weak or no coupling to mutant mtDNA (B, C, E) admit an order-of-magnitude lower increase in wildtype variance as mutant is cleared. Relaxed replication (model A), which combines a target copy number with a strong coupling between mutant and wildtype mtDNA, displays an intermediate increase on wildtype variance as mutant is cleared.

**Figure 6:**
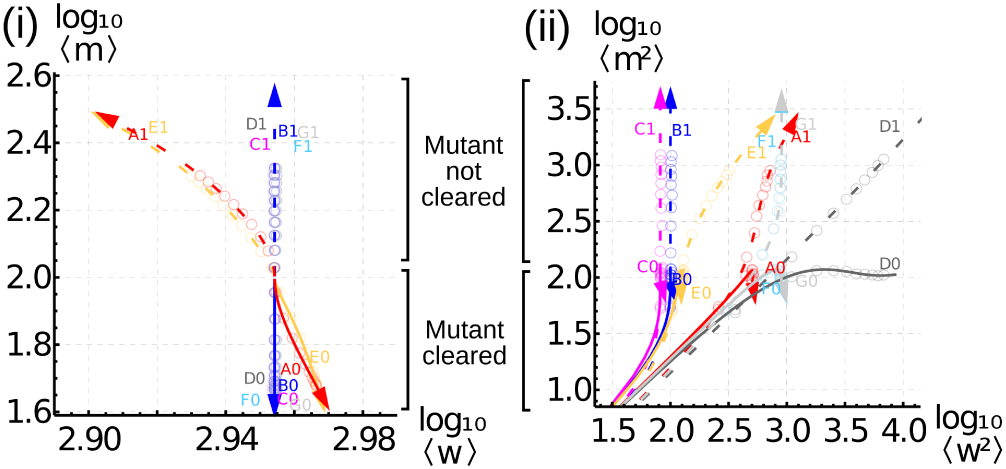
Variability in quality control clearing of mutants from replication errors. Co-evolution of (i) means and (ii) variances of mutant *m* and wildtype *w* mtDNA in cells where replication errors occur. Solid lines (downward trajectories; labels [A-F]0) give the case where quality control is sufficiently strong to clear the resulting mutant molecules (hence decreasing 〈*m*〉 in (i)); dashed lines (upward trajectories; labels [A-F]1) give the case where quality control cannot clear mutants (hence increasing 〈*m*〉). Colours and labels correspond to different control models (see text and Fig. 2). Points show stochastic simulation. Importantly, regardless of the success of quality control in clearing mutants, wildtype mtDNA variability 〈*w*^2^〉 can increase substantially during the action of quality control (rightwards movement in (ii)), and some strategies lead to order-of-magnitude differences in this increase (starred arrow).

Theoretical approaches which only consider the mean behaviour of mtDNA populations (Fig. 6 (i)) cannot account for this cellular heterogeneity, and the important fact that quality control acting to remove mutant mtDNA can also induce variability in wildtype mtDNA (Fig. 6 (ii)). The action of quality control may therefore yield a subset of cells with wildtype mtDNA substantially lower than the mean value across cells – potentially placing a physiological challenge on those cells where wildtype mtDNA is decreased. In addition to the important point that the simple presence of quality control does not guarantee the clearing or stabilisation of mutant load, we thus find that *quality control may have substantial effects on wildtype as well as mutant mtDNA if cellular control couples the two species (Fig. 1 V).*

### Applications III: Variance induced through mutant clearing

The *A* > *G* mutation at position 3243 in human mtDNA is the most common heteroplasmic pathological mtDNA mutation, giving rise to MELAS (mitochondrial encephalomyopathy, lactic acidosis, and strokelike episodes), a multi-system disease. The dynamics of 3243*A* > *G* heteroplasmy are complex and tissue-dependent; its behaviour in blood has been characterised in particular detail using a fluorescent PCR assay for heteroplasmy [38] in a way that allows us to explore our theoretical predictions about mtDNA statistics with mutant clearing.

The authors of Ref. [38] took two blood samples, several years apart, from human patients, and quantified 3243*A* > *G* heteroplasmy (*h*) and mean mtDNA copy number per cell for both samples. While single-cell data is not presented in the publication, progress can be made with the averaged quantities. A strong decrease in (*h*) with time is observed for all patients, confirming that mutant mtDNA is being cleared (Fig. 7(i)). The behaviour of total mtDNA molecules per cell (*w* + *m*) is less consistent, with a range of large increases and moderate decreases in total number. As shown in Fig. 7(ii)-(iii), patient-to-patient variance in both *w* and *m* increases with time in conjunction with *h* decreases.

**Figure 7:**
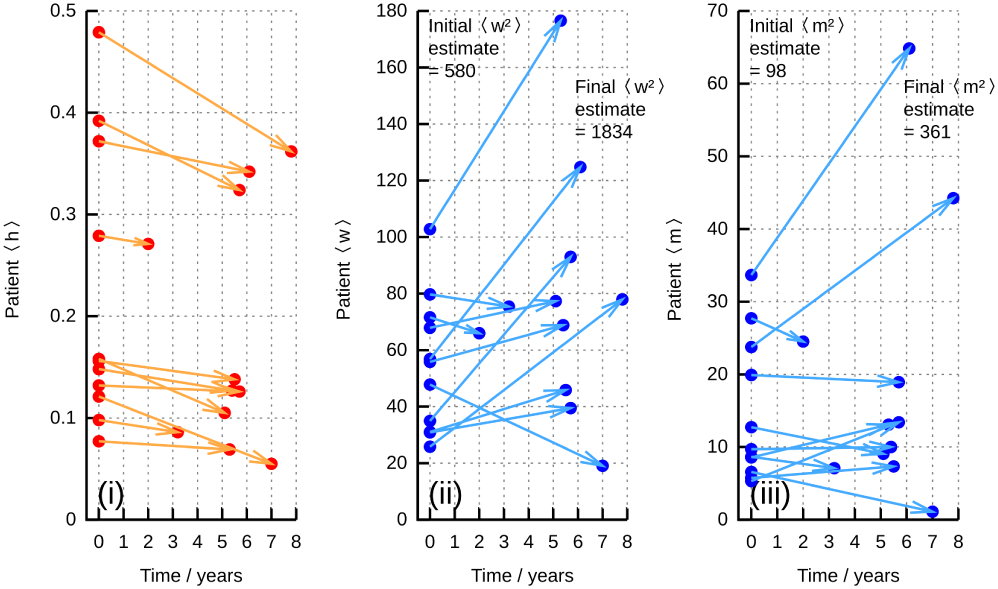
MtDNA evolution in MELAS patients from Ref. [38]. (i) Changes in mean heteroplasmy 〈*h*〉 between two samples from patient blood, showing a general decrease in 〈*h*〉 over time. (ii) Changes in mean cellular wildtype content *w* between these samples. (iii) Changes in mean cellular mutant content *m*. In both (ii) and (iii), an overall increase in variance over time is observed: hence 〈*w*^2^〉 and 〈*m*^2^〉 increase while 〈*h*〉 decreases, consistent with Fig. 6.

Although the measurements in Ref. [38] are averages over groups of cells, an approximate quantitative comparison of these data with the predictions of our theory can be made. In the spirit of ‘back-of-the-envelope’ calculations [39], we estimate that each sample of cells giving rise to a measurement corresponds to approximately 10^3^ cells (see SI Section 8). Then, using an estimate of *τ* = 5 days (by comparison with other mammalian species, as above), we find that a selective pressure of ∊_8_ ≃ 1.2 × 10^−4^ day^−1^ matches the observed decrease in heteroplasmy, corresponding, for example, to a decrease from 0.15 to 0.14 over 8 years, as in Fig. 7(i). Solving the ODEs resulting from this system, using control model D as the simplest case, predicts increases in cell-to-cell copy number variance of approximate magnitude 〈*w*^2^〉 ~ 10^5^ and 〈*m*^2^〉 ~ 2 × 10^4^ over 8 years. We can translate these cell-to-cell values into the variance expected across samples of cells by dividing by the number of samples (taken as 10^3^ as above). The resulting variance corresponds, for example, to expected standard deviations in wildtype and mutant copy number after 8 years of 10.0 and 4.0 respectively for a sample with 〈*w*〉 = 85 and 〈*m*〉 = 15 (consistent with the increasing spread of values in Fig. 7(ii)-(iii)). We used the Kolmogorov-Smirnov test respectively to test the alternative hypotheses that the experimentally-observed 〈*w*〉 and 〈*m*〉 at the later time point differed from those predicted by our model; no test yielded *p* < 0.05 (see SI Section 8). Of course, an absence of support for an alternative hypothesis cannot be taken as support for a null hypothesis, but shows that the existing experimental data is not incompatible with our model.

The observations in Fig. 7 support the predictions made in Fig. 6, where mutant load is decreased but variance in wildtype copy number (and mutant copy number) increases. The bottom-left quadrant of Fig. 6 (ii) shows that different control mechanisms display similar initial behaviour; follow-up studies on these patients could be used to distinguish possible mechanisms for mtDNA control. For example, if the rate of wildtype variance increase decreases over time, models B, C, and E are more likely; if wildtype variance continues to increase, models A, D, F, and G and more likely. More detailed model discrimination based on the time behaviour of *w* and *m* variances are possible (Fig. 2), and can be performed using statistical methods accounting for mean and variance behaviour [29, 40].

## Discussion

A general, bottom-up theory has been produced to describe the time behaviour of cell-to-cell variance in mtDNA populations subject to controlled biogenesis and/or degradation, mutation, selection, and cell divisions. This theory is based around the microscopic behaviour of mtDNA molecules, allowing a hitherto absent connection between widely-used ‘effective’ statistical genetics approaches (Eqn. 13) and measurable biological quantities, and motivating experiments to further elucidate the mechanisms acting to control mtDNA (described in SI Section 7). We have shown that the predictions of this theory agree with experimental observations of mixed mtDNA populations, and that the application of appropriately validated mathematical theory allows us to make estimates of important biological quantities that remain challenging to directly address with experiments. Our theory describes the cell-to-cell variability in mtDNA populations and thus provides a framework with which to understand the inheritance and onset of mtDNA diseases [6, 14].

Our theoretical platform unifies several existing modelling approaches that have driven advances in the study of mtDNA populations. We have specifically demonstrated that the ‘relaxed replication’ model [19, 1, 33] (our model A), simple birth-death models [14, 20] (our model D), and cellular controls based on homeostatic principles [15, 16] (our models B, C, G) can naturally be represented within our framework. As a result, analytic expressions for the expected behaviour of heteroplasmy variance and other population statistics can readily be extracted for these and other mtDNA models (see SI Section 3), allowing the detailed characterisation of mtDNA dynamics, including the probability of crossing disease thresholds [7], which can be computed from heteroplasmy statistics [14]. We have also used the theoretical ideas developed herein to refine a widely-used model for mtDNA populations of changing size (the Wright formula), explicitly connecting it with cellular processes and allowing a link between physical and genetic quantities (Eqn. 14, Fig. 5).

Further, our theory also describes the dynamics of heteroplasmy change with time in the presence of selective pressure for one mtDNA type. Given an initial heteroplasmy *h*_0_ and a selective pressure *β* (positive *β* corresponding to positive selection for the mutant mtDNA type), we find (see SI Section 5) that heteroplasmy evolves according to:

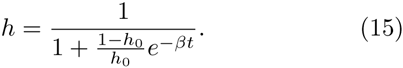

This behaviour immediately motivates a transformation allowing the evolution of heteroplasmy to be compared across different starting values *h*_0_:

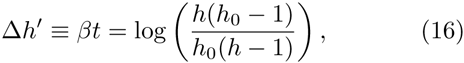

allowing, as in our previous work [23], heteroplasmy results from different biological samples to be compared together, accounting for different initial heteroplasmies (in other words, the same selective pressure will produce the same Δ*h*′ regardless of *h*_0_).

It is likely that control mechanisms found in biology have nonlinear forms (for example, sigmoidal response curves are common in cellular signalling). We have shown that a linearisation satisfactorily describes some non-equilibrium behaviour (for example, in the case of our cell division model) but further investigation of more general nonlinear behaviour, and modulation of wider cell behaviour by mtDNA populations (for example, by influencing cell cycle progression [15]), are important future developments. In SI Section 1 we discuss a linear stability analysis of our expressions for mean mtDNA behaviour, which highlights a link between the ‘sensing’ of an mtDNA species (in the sense that the presence of that species modulates replication or degradation rates) and the ability to control the mean level of that species.

The control of stochastic systems is a well established field within control theory [41]. Optimal control mechanisms addressing the mean and variance of stochastic processes have been derived in a variety of contexts (see, for example, [42] and citations therein), particularly in financial applications [43], and often find tradeoffs between controlling the mean and variance of a process. We observe a comparable tradeoff, that tight control on moments of one species leads to loose control on another. We have focussed on providing a general theoretical formalism with which to treat any given control mechanism; it is anticipated that the above treatment may also be of value in describing heterogeneity in other systems where replication and/or death rates of individuals depend on feedback from current numbers of individuals (for example, through terms describing competition for resources in ecology). Within the context of mtDNA populations, we anticipate that this theoretical framework will assist in understanding natural processes of mtDNA inheritance and evolution within an organismal lifetime (including segregation and increasing variance with age) [14, 23], and informing applied approaches to control mitochondrial behaviour with genetic tools [4, 5].

## Appendix

### A1. Interpretation of relaxed replication

The relaxed replication model [19, 1] describes a cellular population of mtDNA molecules according to the following algorithm. MtDNAs randomly degrade as a Poisson process with rate 1/*τ*. Every timestep Δ*t*, the value of *C*(*w, m*), a deterministic function of *w* and *m*, is computed, then Δ*tC*(*w, m*) mtDNAs are added to the population. The *genetic* properties of these added mtDNAs are random – each is assigned a genetic type based on a random sampling of the existing populations – but their *physical* properties (i.e. the total copy number added at each step) is deterministic. We argue that, as both replication and degradation of mtDNAs depend on complicated behaviour and thermal, microscopic interactions, it makes more sense to model both processes as stochastic. Thus, *C*(*w, m*) is interpreted as the rate of a Poisson process describing replication, just as 1/*τ* is the rate of a Poisson process describing degradation. This interpretation reconciles the nature of the two processes.

Although this feature is less interesting than the underlying scientific behaviour, the original algorithm also raises a (not insurmountable) technical problem with implementation. If a timestep Δ*t* < 1/*C*(*w, m*) is chosen, the algorithm will never add any mtDNA to the system. But, in order to suitably characterise the stochastic degradation without using the Gillespie algorithm [44], it is desirable to choose a timestep Δ*t* as low as possible. There is therefore a risk that one or other of the deterministic replication and stochastic degradation processes is inadequately captured in a given simulation protocol.

We have illustrated the excellent agreement between our theoretical approaches and stochastic simulation (for example, Fig. 2). To quantitatively connect with previous analyses of specific control strategies, we confirm that the behaviour for model A (relaxed replication) matches that observed in previous simulation studies [19] with a back-of-the-envelope calculation [39]. The rate of variance increase with time with *τ* = 5 (comparable in magnitude to the (1 – 10) × ln 2 days used in Ref. [19]) and w_*opt*_ = 1000 from Fig. 2 is roughly 40 day^−1^. Considering 50 years of evolution of this system, we expect a standard deviation of roughly 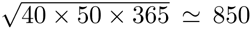 in mutant copy number. This value is consistent with the simulations in Ref. [19].

We connect to an additional numerical result in Ref. [1]. In the absence of a mutant population, the variance of the wildtype population was reported to be stable at *w*_*opt*_/(2*α*), with the original model interpretation of mtDNA replication as deterministic. Under the interpretation of stochastic replication, an absent mutant population (*m_ss_* = 0) permits stability in the wildtype population variance, which after a little algebra is calculated to be *w_opt_*/*α*. Intuitively, modelling both replication and degradation as stochastic does not affect mean copy number but does increase variance.

### A2. Interpretation of Wright formula for mtDNA

The Wright formula

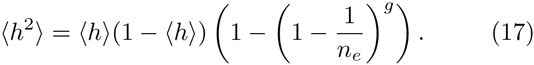

has been proposed as a model for the time evolution of heteroplasmy variance 〈*h*^2^〉 in a population with effective size *n_e_* subject to random partitioning at each of *g* generations. This picture has been successfully employed to investigate heteroplasmy distributions in real systems [22], with *n_e_* and *g* interpreted as parameters of the theory without immediate biological interpretation.

The mapping of the original genetic system considered by Wright [21] to cellular populations of mtDNA requires some discussion. If ‘generations’ are interpreted as cell divisions, the mechanism by which mtDNA copy number is redoubled between divisions is assumed by the model to be deterministic. Cell divisions will result in a halving of the mtDNA population. Application of the Wright model assumes that the original population is thenceforth recovered with no increased variance in the population. In other words, the mtDNA population is assumed to exactly double between divisions with no stochasticity in the process. As we underline in the Main Text, the effects of (inevitable) stochasticity due to mtDNA turnover are not explicitly captured by the Wright formula. Other complications exist, as described in [22], but play less important roles here. As a result, the ‘bottleneck size’ *n_e_* cannot immediately be interpreted as an observable minimum cellular copy number of mtDNA molecules (a quantity that is reported by, for example, a qPCR experiment measuring cellular mtDNA content), but rather the size of an effective ‘founder’ population.

## Supplementary Information

### S1. Expansion of the control law

If we write *∊*_*w*_ = *w* − *w*_*ss*_, ∊_*m*_ = *m* – *m_ss_*, the expansion of replication and degradation rates λ(*w, m*) and *ν* (*w,m*) about the steady state {*w_ss_, m_ss_*} gives

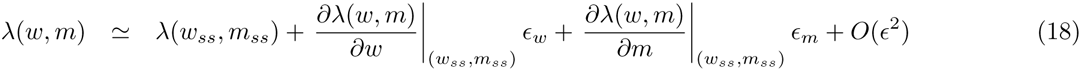

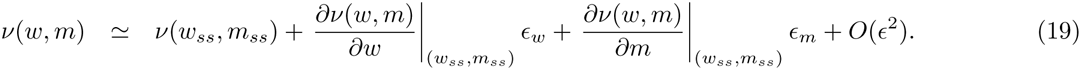

This expansion represents a model of a given control strategy λ(*w, m*), *ν*(*w,m*), which, if the original function is well behaved, we expect to reasonably reflect behaviour of the system close to {*w_ss_*, *m_ss_*}. Simulation results show that this expectation is fulfilled for a wide variety of cases (see figures in the Main Text).

To find *w_ss_* and *m_ss_* we solve the equations describing the deterministic behaviour of the system:

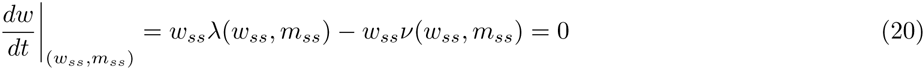

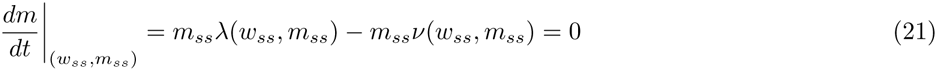

It will be observed that for this steady state to exist, the terms *λ*(*w_ss_*, *m_ss_*) and *ν*(*w*_*ss*_, *m*_*ss*_) in Eqns. 18-19 must be equal. We can write the general expansion form of *λ*(*w, m*) and *ν*(*w, m*), truncated to first order, as

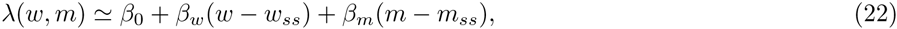

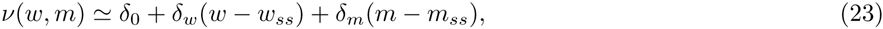

with **β*_w_* = ∂λ/∂*w*|_*w_ss_,m_ss_*_, *β*_*m*_ = ∂λ/ ∂*m*|_*w_ss_,m_ss_*_, *β*_*m*_ = ∂ν/∂*w*|_*w_ss_,m_ss_*_, δ_*m*_ = ∂ν/∂*m*|*w_ss_,m_ss_*. Clearly, to support convergence to a steady state, *β*_*w*_ and *β*_*w*_ must be negative and δ_*w*_ and δ_*m*_ must be positive. Given this model for control dynamics, we next characterise the variance of the system. We can thus describe the system with a set of *R* = 4 processes with rates

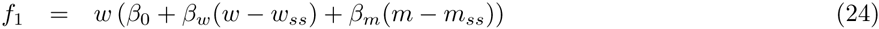

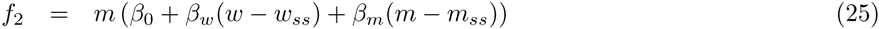

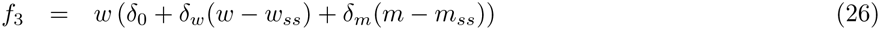

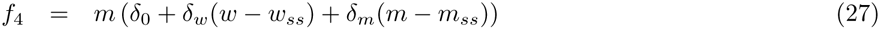

and stoichiometry matrix describing the effects of these reactions on the *N* = 2 species we consider as

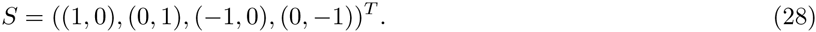

Using index *i* = 1 to correspond to species *w* and *i* = 2 to correspond to species *m*, the master equation for the system, describing the time evolution of *P*_*w,m*_ (the probability of observing *w* wildtype and *m* mutant mtDNAs) can then be written

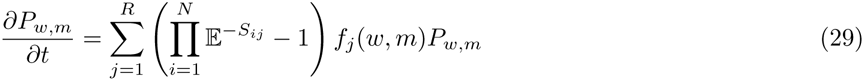

where 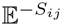 takes its normal meaning as a raising and lowering operator [24], adding *−S_ij_* to each occurrence of index i that follows it on the right (so e.g. as *S*_11_ = 1 and *w* corresponds to index 1, 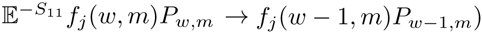.

The potential nonlinearities and coupling between species in this equation prevents a full general solution. To make progress, we employ Van Kampen’s system size expansion [24, 25] and write *w* = *ϕ*_*w*_Ω + ξ_*w*_Ω^1/2^, *m* = *ϕ*_*m*_Ω+ξ_*m*_Ω^1/2^, representing copy numbers as the sum of deterministic components *ϕ_i_* and fluctuation components ξ_*i*_ scaled by powers of system size Ω. Following the standard expansion procedure, by writing 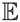, *P_w,m_* and *f_i_* in terms of Ω and collecting powers of Ω in Eqn. 29, first gives equations for the deterministic components of the system (corresponding straightforwardly to the macroscopic rate equations):

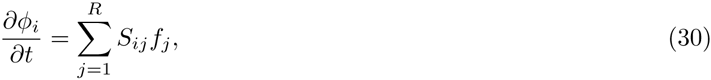

then gives a Fokker-Planck equation for the time behaviour of the fluctuation components in terms of the bivariate probability distribution Π(ξ, *t*) of ξ = (ξ_*w*_, ξ_*m*_) at time *t*:

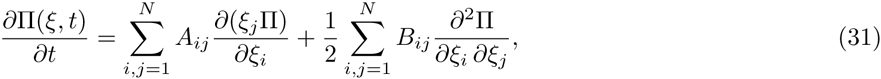

where

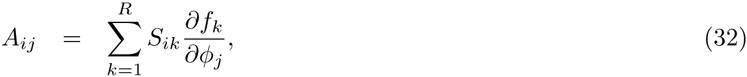

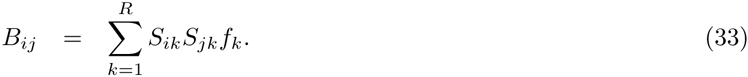

The form of Eqns. 24-27 and Eqn. 28 gives, for steady state copy numbers and *δ*_0_ = *β*_0_, *A*_11_ = *κ*_*w*_*w*_*ss*_, *A*_12_ = *κ*_*m*_*w*_*ss*_, *A*_21_ = *κ*_*w*_*m*_*ss*_, *A*_22_ = *κ*_*m*_*m*_*ss*_, *B*_11_ = 2*β*_0_*w*_*ss*_, *B*_22_ = 2*β*_0_*m*_*ss*_, *B*_12_ = *B*_21_ = 0, where *κ*_*w*_ = (*β*_*w*_ − *δ*_*w*_), *κ*_*m*_ = (*β*_*m*_ – *δ*_*m*_). From this Fokker-Planck equation expressions for the moments of *ξ_i_* can be extracted [24], leading to the expressions:

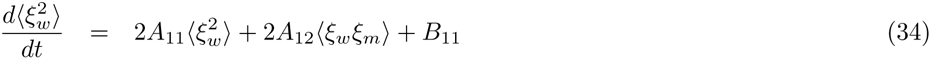

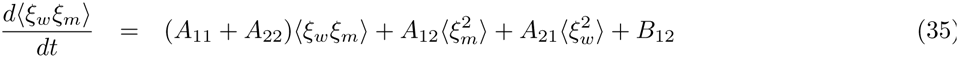

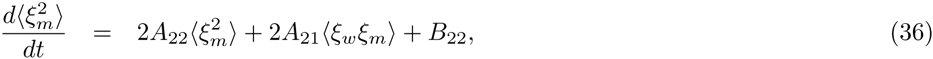

A linear stability analysis of the deterministic ODEs describing mean behaviour is straightforward to perform. Linearising Eqns. 20-21 about (*w*_*ss*_, *m*_*ss*_) gives

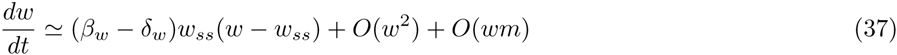

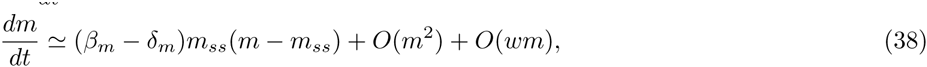

from which it is straightforward to see that if *κ*_*w*_ < 0 and *κ*_*m*_ < 0, the mean dynamics of *w* and *m* respectively are linearly stable. This condition is met for *w* and *m* in control laws A and E, for *w* in B, C, and F, and for neither in D. These specific examples illustrate the principle that if a species is explicitly ‘sensed’ – in the sense that it modulates replication or degradation rate – its mean dynamics can be controlled to be linearly stable. If a species is not explicitly sensed (replication and degradation are not functions of its copy number) then its mean dynamics are not explicitly linearly stable, but may be ‘balanced’, as the corresponding *κ* term is zero. Perturbations in these unsensed ‘balanced’ variables are neither damped away by control nor guaranteed to explode with time; hence the variables are unconstrained but not explicitly unstable.

### S2. Full solutions for steady state ODEs

Eqns. 34-36 can be solved exactly for the *A_ij_*, *B_ij_* corresponding to the steady state condition above. The complete solutions for arbitrary 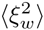, (ξ_*w*_ξ_*m*_), 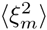 at *t* = 0 are lengthy and do not allow much intuitive interpretation. For 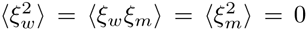 at *t* = 0 (noiseless initial conditions), we can solve and separate the long-term behaviour from the transient behaviour. The transients, given by Eqns. 42-44 below, involve terms in *t*′ ≡ exp((κ_*m*_*m*_*ss*_ + κ_*w*_*w*_*ss*_)*t*). As the κ_*i*_ are nonpositive (for stability, *β*_*i*_ are nonpositive and *δ_i_* are nonnegative), *t*′ is either a constant or an exponentially decaying function of time *t*; in the cases we consider, it either decays with time (all models except D) or the associated term is always zero (model D).

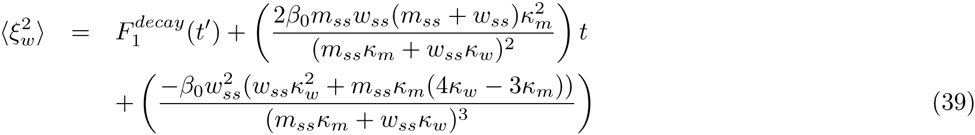

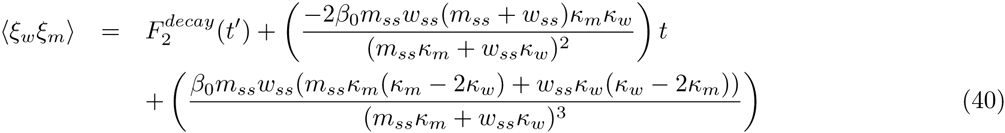

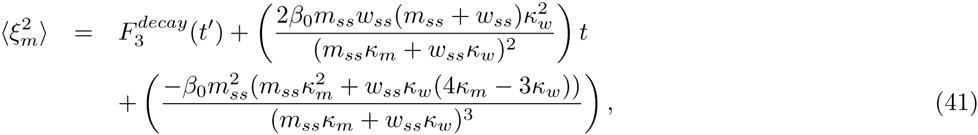

where the 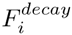 functions characterising the transient behaviour are

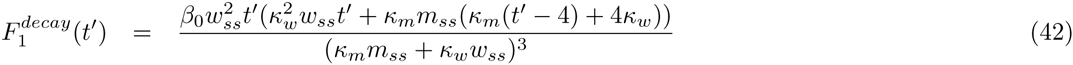

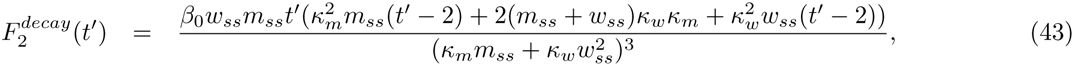

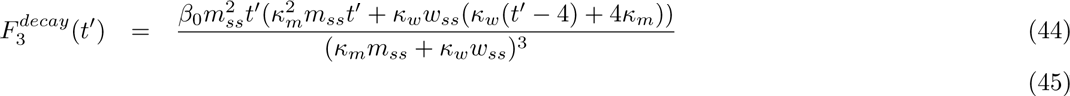

Eqns. 39-41 with Eqns. 42-44 then give the full transient behaviour displayed in Fig. 2, decaying to the aforementioned linear behaviour with characteristic timescale (*κ*_*m*_*m*_*ss*_ + *κ*_*w*_*w*_*ss*_).

### S3. Interpretation and behaviour of specific control strategies

The mechanisms described in the Main Text have simple interpretations in the language of stochastic processes. Mechanism A, as discussed, corresponds to the ‘relaxed replication’ picture studied previously (Appendix A1). Mechanism D corresponds to independent birth-death processes acting on both species (analysed in [14]). Mechanism F corresponds to an immigration-death process acting on the wildtype (the dependence of replication rate on 1/*w* means that overall production is constant with time), and model C can be regarded as a birth-immigration-death process on the wildtype (analysed in [20]). Both these mechanisms are thus expected to tightly control wildtype behaviour (including controlling variance: immigration-death processes yield a constant steady-state variance), but do not sense (and, therefore, do not apply feedback to) mutant load. We also note that mechanisms F and G are in a sense ‘dual’, in that they apply similar control manifest through replication with rate ∼ 1/*w* and degradation with rate *w* respectively. The previous result that control applied to (a) biogenesis rates and (b) degradation rates yields similar population behaviour is visible in the behaviour of F and G in Fig. 2.

Table 1 gives, for each example control strategy in the Main Text, the corresponding steady state {*w_ss_*, *m_ss_*} and the expansion terms for the strategy **β*, δ*. The corresponding behaviours of variances, from Eqns. 39-41, are given in Table 2.

**Table 1:**
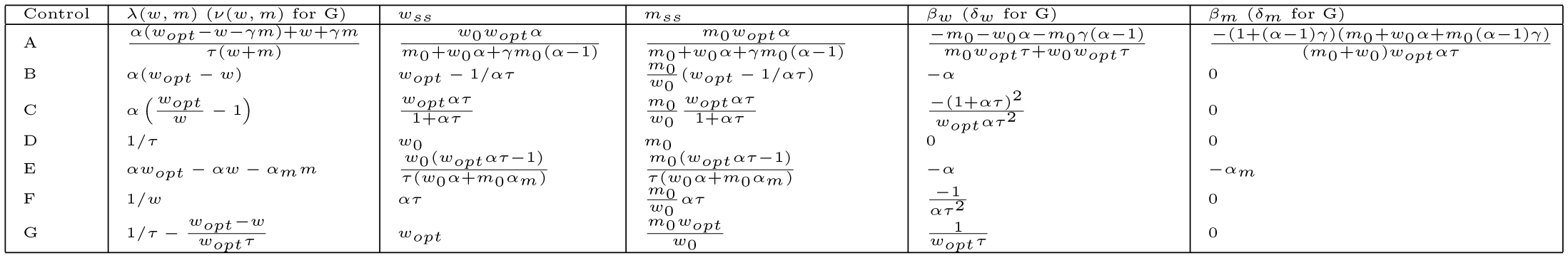
Steady states and expansion terms for control strategies A-G.

**Table 2:**
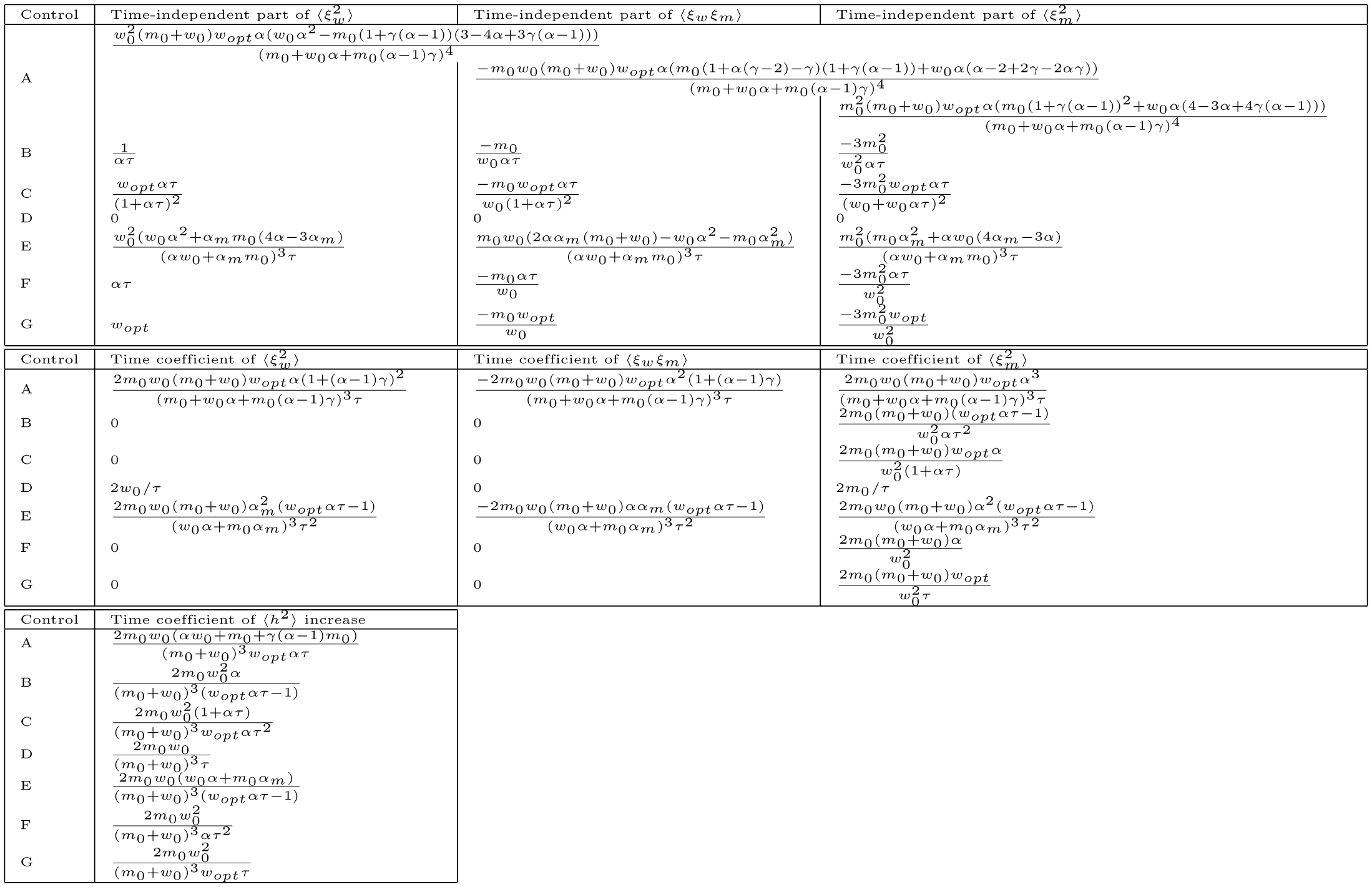
Post-transient time behaviour of copy number and heteroplasmy variances in control strategies A-G.

In the Main Text we discuss the implications of an increasing extinction probability of one mtDNA type. A nonnegligible extinction probability challenges the validity of the linear noise approximation and leads to departure from results derived using the system size expansion. To illustrate this behaviour, we reduce the characteristic timescale of the parameterisations used to explore models A-G in the Main Text, setting *τ* = 1 rather than *τ* = 5, and simulate for a longer time window (see Fig. 8). It will be observed that as extinction probability increases (as 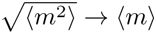), the numerical behaviour departs from that predicted analytically; in particular, the increase of 〈*h*^2^〉 slows from a linear to sublinear regime as discussed in the Main Text.

**Figure 8:**
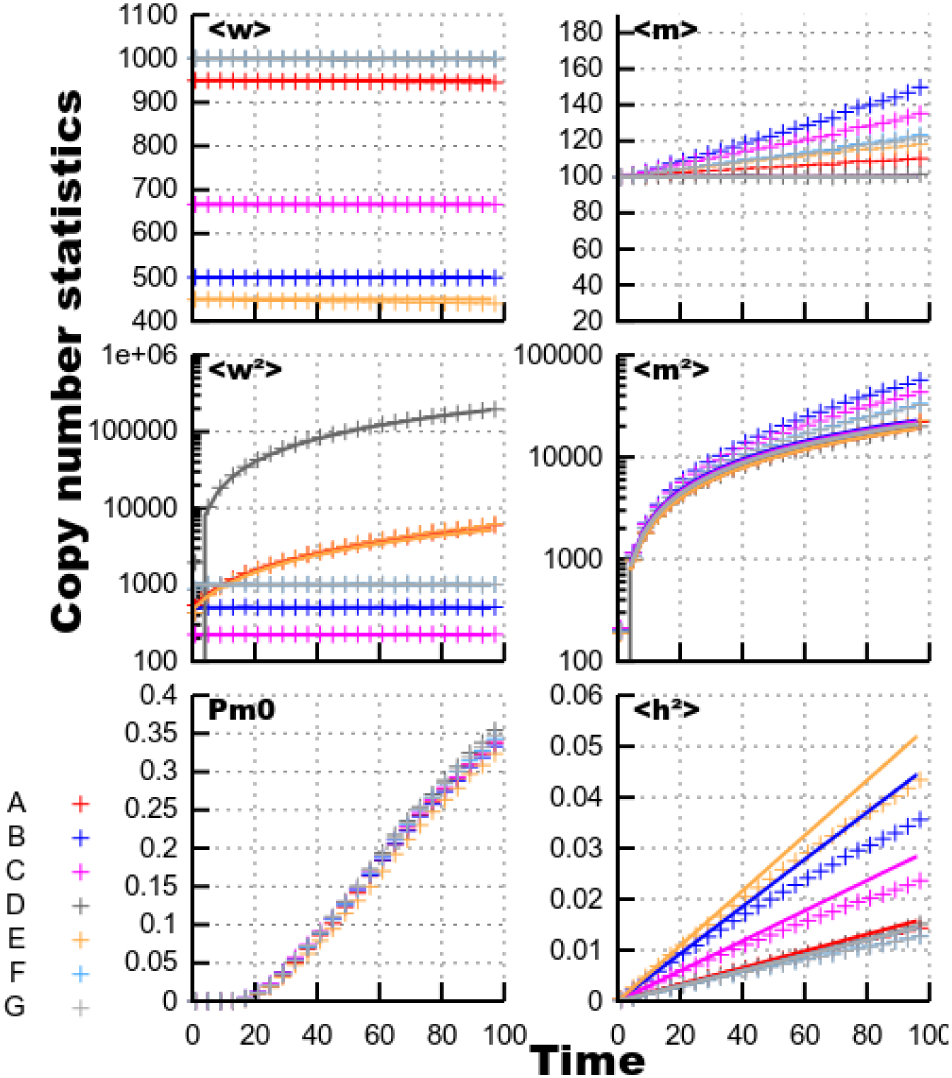
Influence of fixation on expansion analysis. The models from the Main Text, simulated for a longer time window and for a shorter characteristic timescale *τ*, illustrating the behaviour of the systems when extinction becomes possible. *Pm*0 gives the numerically computed probability that *m* = 0; it can be seen that an increase in this quantity corresponds to a moderate increase of 〈*m*〉 and 〈*m*^2^〉 relative to their predicted values, and a decrease of 〈*h*^2^〉 relative to its predicted value (shifting towards a sublinear increase as discussed in the Main Text).

We note that, in some physiological circumstances, the representation of one mtDNA type in a cellular population may be low – for example, the appearance of one mutant mtDNA through *de novo* mutation or replication error, or the presence of a small percentage of a foreign mtDNA haplotype due to carryover in gene therapies [6]. In these cases, a non-negligible extinction probability may occur quickly and the transition of 〈*h*^2^〉 to a sublinear, or flat, regime will be an important aspect of the long-term dynamics. In the case of mtDNA disease inheritance, however, situations with a macroscopic fraction of mutant mtDNA are often the most important, due to the presence of a ‘heteroplasmy threshold’ [7] beyond which disease symptoms manifest. With two mtDNA haplotypes represented in comparable proportions in the cell, our linear analysis holds and can be used to describe heteroplasmy variance in somatic and germline cells.

### S4. Steady state solution

Eqns. 34-36 give, for steady state,

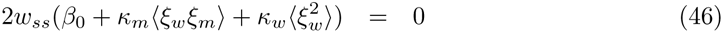

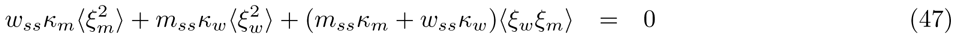

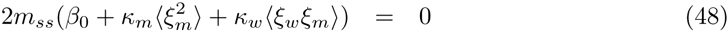

Attempting to solve these equations for 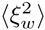, then (ξ_*w*_ξ_*m*_), then 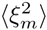 first gives 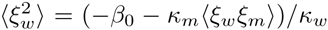, then 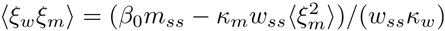, leaving Eqn. 47 reduced to

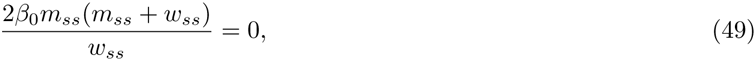

a condition only fulfilled (due to the non-negativity of *m_ss_* and *w*_*ss*_) if *m_ss_* = 0. If one proceeds through the analysis by first solving for 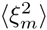, then 〈ξ_*w*_ξ_*m*_〉, then 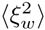, a symmetric expression is obtained

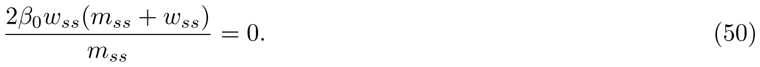

Eqns. 49-50 illustrate the symmetry in the system: if the copy number of either species is zero then a situation where variance does not increase is supported (but not inevitable: compare the behaviour of relaxed replication model (A, fixed variance) and the birth-death model (D, increasing variance) in the case of zero mutant population).

The effect of selection, mutations, and replicative errors on mtDNA variances can straightforwardly be included in this analysis. In this general case, we replace Eqns. 1-4 in the Main Text with:

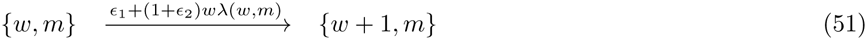

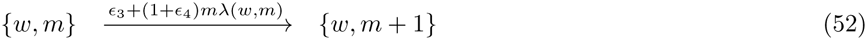

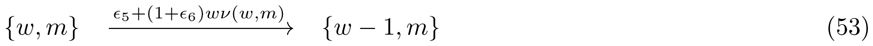

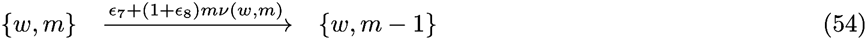

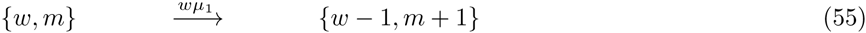

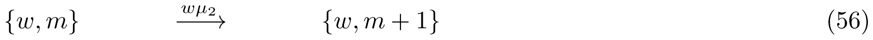

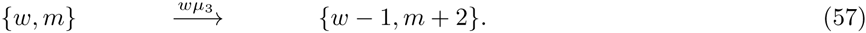

Here, we have added processes corresponding to spontaneous mutation of a given wildtype mtDNA (*μ*_1_), and two types of replicative error affecting wildtype mtDNA, giving rise to one (*μ*_2_; original molecule remains intact, new molecule is mutated) and two (*μ*_3_; both original and new molecules are mutated) mutant mtDNAs respectively. Differences manifest either through replicative or degradation advantages (or both) are incorporated with even-indexed *∊_i_* (providing multiplicative changes to the bare rates) and odd-indexed *∊_i_* (providing additive changes). We thus have two ways of provoking selective advantages in each case: increasing wildtype biogenesis, increasing mutant biogenesis, increasing wildtype degradation, and increasing mutant degradation.

Fig. 9 shows example trajectories arising from each of our control mechanisms in the presence of the mutation processes above, and the selective pressures (*∊*_3_, *∊*_4_, *∊*_5_, *∊*_6_) that favour mutant mtDNA. An excellent agreement between ODE theory and stochastic simulation is again illustrated, and there is substantial similarity between the behaviours caused by selection (favouring mutant mtDNA) and mutation (producing mutant mtDNA). In several cases (mechanisms B, C, F, G), 〈*m*〉 and 〈*m*^2^〉 simply increase exponentially with time under favourable selective or mutational pressures; this situation straightforwardly gives rise to a sigmoidal change in heteroplasmy 〈*h*〉 ∼ 1/(1 + e^−Δ*ft*^(1 – *h*_0_)/*h*_0_), with Δ*f* an effective selective difference, as used in previous work [45, 23]. In mechanisms coupling wildtype and mutant content (A and E), mutant increase is slower and accompanied by a decrease in wildtype, attempting to keep total copy number constant. In these circumstances, variance behaviour can be more complex: for example, under relaxed replication with pressure favouring mutant mtDNA, 〈*w*^2^〉 initially increases then subsequently decreases as 〈*w*〉 decreases in magnitude. Mechanism D, where control does not couple mutant and wildtype, has correspondingly perpendicular trajectories in (〈*w*〉, 〈*m*〉) space under different selective pressures, but the coupling action of the mutation operations lead to curved trajectories under mutational influence.

**Figure 9:**
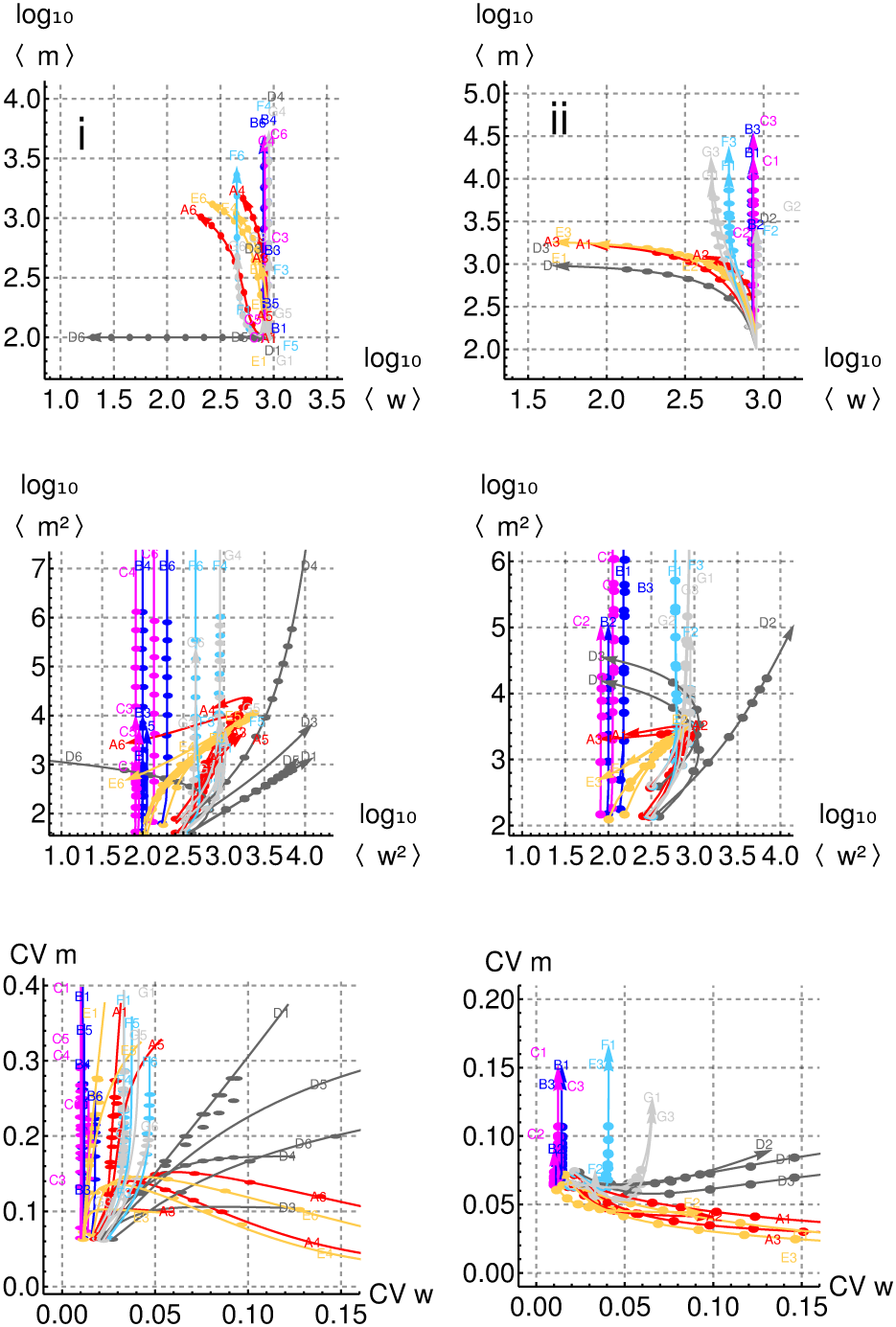
MtDNA copy number and variability under mutational and selective advantages for mutant mtDNA. Mean, variance, and CV trajectories with (i) selection pressures, and (ii) mutation rates favouring mutant mtDNA, under models A-G from the text. Some control strategies (A, E) keep mutant relatively bound but sacrifice wildtype and provoke large increases in variability; others (B, C, F, G) focus on wildtype stability, allowing mutant to grow unbound. Labels give the control model (letter) and the parameter varied (∊_3_ = 20; ∊_4_ = 1; ∊_5_ = 20; ∊_6_ = 1 for (i); *μ*_1_ = 0.1; *μ*_2_ = 0.1; *μ*_3_ = 0.1 for (ii)); all other ∊, *μ* parameters are set to zero. Results are shown for theory (lines) and stochastic simulation (points), progressing from an initial condition with *w*_0_ = 900, *m*_0_ = 100 with the parameterisations in Fig. 2.

### S5. Heteroplasmy

For a general function *h* = *h*(*x,y*),

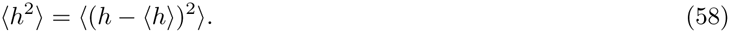

We will consider an expansion about (*x*_0_, *y*_0_), a state such that *h*(*x*_0_, *y*_0_) = 〈*h*〉. Using the first-order Taylor expansion of *h*(*x, y*) around (*x*_0_, *y*_0_):

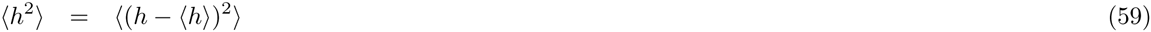

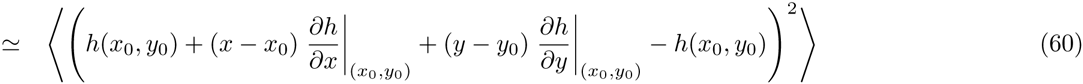

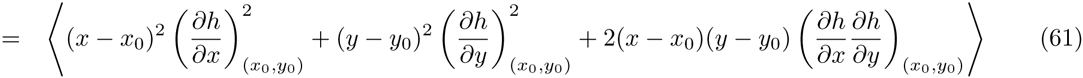

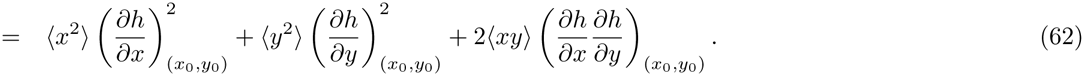

We now consider *h*(*x, y*) = *x/y*, so that

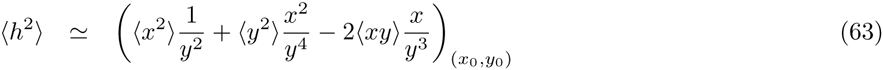

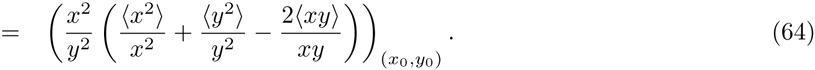

Finally, given that *x*_0_ = 〈*x*〉 and *y*_0_ = 〈*y*〉, and setting *x* ≡ *m* and *y* ≡ *w* + *m*, we obtain

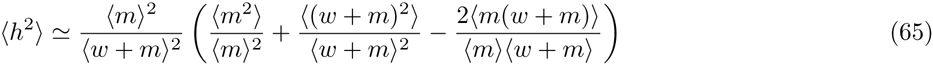

To see that exponential growth or decay in one mtDNA type while the other remains constant gives rise to sigmoidal heteroplasmy dynamics, consider (without loss of generality) *m* = *h*_0_*n*_0_*e*^*βt*^, *w* = (1 – *h*_0_)*n*_0_, where *n*_0_ is an initial population size which will cancel. Then, as *m* (and hence *n* = *m* + *w*) increases with time,

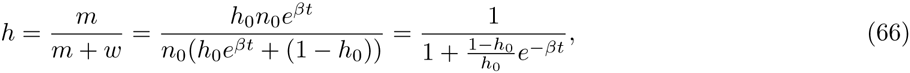

as used in Refs. [6] and [14], with *β* corresponding to a selective pressure (in this derivation, positive *β* favours mutant mtDNA).

### S6. Fokker-Planck terms for nonequilibrium regimes

The system size expansion approach above can be applied to the general system without employing an expansion of the control strategy about a steady state, by considering the processes

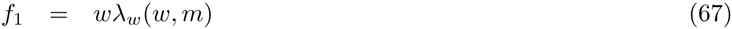

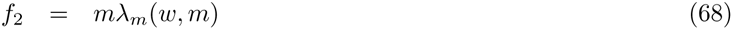

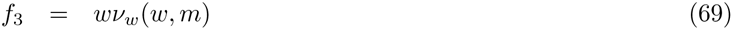

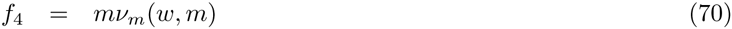

If the expansion about steady state is not used, the corresponding terms are

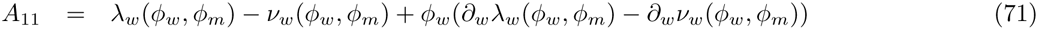

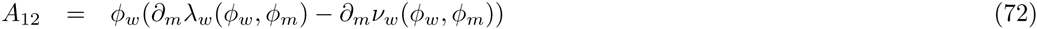

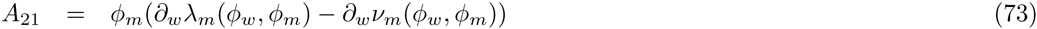

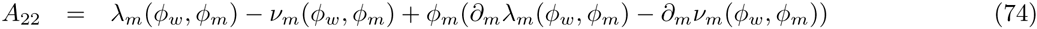

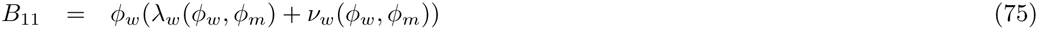

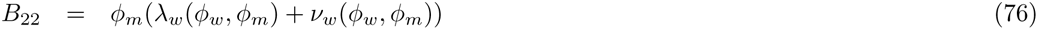

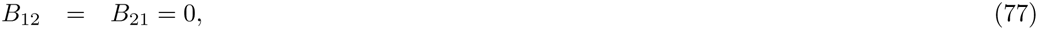

where ∂_*x*_*f*(*ϕ*_*i*_; *ϕ*_*j*_) means 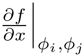. We include the mutational processes in the text by adding *f*_5_ = *μ*_1_*w*, *f*_6_ = *μ*_2_*w*, *f*_7_ = *μ*_3_*w* and setting the corresponding stoichiometry matrix to

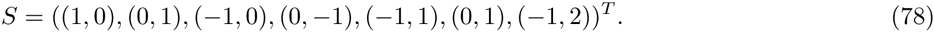

If λ_*w*_ = λ_*m*_ = λ and *ν*_*w*_ = *ν*_*m*_ = *ν* (no selective differences between mtDNA types), the Fokker-Planck terms

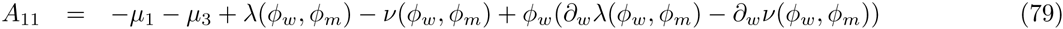

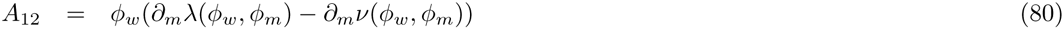

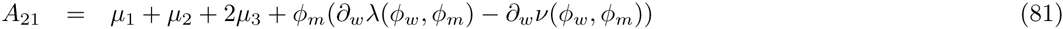

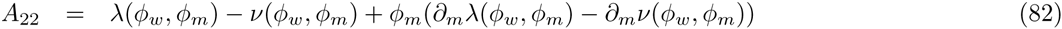

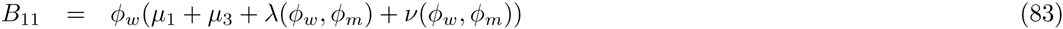

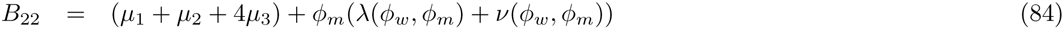

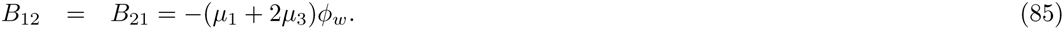

Including selection terms (without mutation) requires no change to the original structure of reactions and stoichiometries and immediately gives

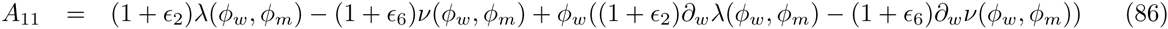

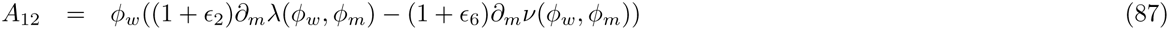

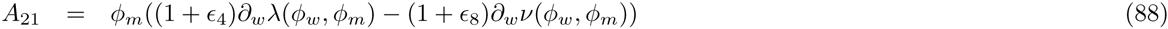

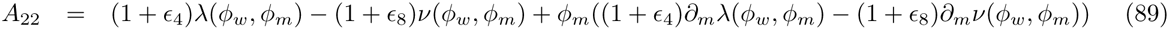

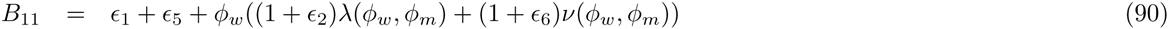

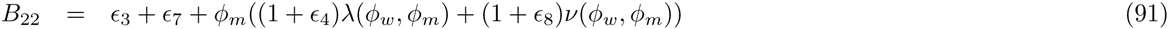

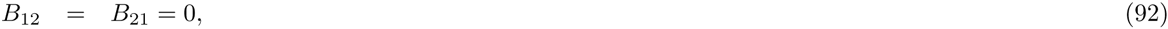

The same approach as above can be used to obtain Eqns. 34-36 for the time evolution of fluctuation moments, this time valid for a full temporal trajectory of the system.

### S7. Experimental observations to distinguish mechanisms

Our theoretical results suggest measurements to further elucidate the control mechanisms underlying mtDNA evolution within cells, without using heteroplasmy variance 〈*h*^2^〉 (the shortcomings of which are manifest because seven different feedback controls all yield the same dynamics in 〈*h*^2^〉′ – Fig. 2), and in conjunction with further molecular elucidation of processes governing mtDNA [46, 2] which providing bounds on the types and rates of molecular processes involved (for example, disallowing unphysically high rates of mtDNA replication).

If 〈*w*^2^〉 increases with time, mechanisms with weaker constraints on wildtype copy number are more likely (including relaxed replication (A), mechanisms sensing a combination of mutant and wildtype copy number (E), and the case with no feedback (D)). If 〈*w*^2^〉 is low and constant, mechanisms involving differential (B) or ratiometric (C) control are likely. If 〈*w*^2^〉 is high and constant (of the order of 〈*w*〉), mechanisms resembling immigration-death processes (with propagation scaled by inverse copy number, F and G) are more likely. The behaviour of 〈*wm*) can be used to further distinguish mechanisms which strongly couple wildtype and mutant (including relaxed replication and total copy number control) from those with less coupling.

In all these cases, the likelihood functions associated with specific biological observations will be complicated. Model selection and inference in this case could be performed through comparison to simulation, or using likelihood-free inference [47] for the mean and variance of mtDNA populations [40].

### S8. Back-of-the-envelope calculations for leukocyte heteroplasmy measurements

Average cellular mtDNA copy number measurements in Ref. [38] are made by normalising the signal from the mtDNA-encoded *ND1* gene by that from the nuclear-encoded *GADPH* genes using real-time PCR using iQ Sybr Green on the BioRad ICycler. The published protocol [48] for this technique suggests using 50ng-5pg of genomic DNA. Diploid human cells contain ∼ 6pg of genomic DNA; the mass of several hundred (much smaller) mtDNA genomes is negligible by comparison. The protocol thus implies the presence of 1-10000 cells’ genomic DNA content; we assume 1000 as an estimate consistent with qPCR standards (Joerg Burgstaller, personal communication).

In our analysis of the data from Ref. [38] we use *τ* = 5 days and the processes:

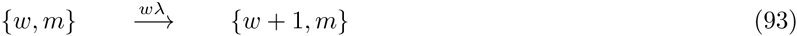

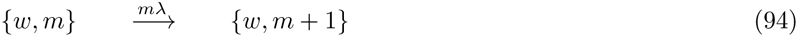

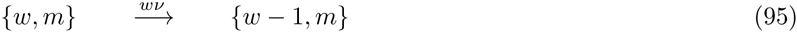

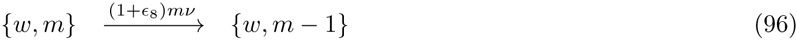

with λ = *ν* = 1/*τ*, and ∊_8_ a selective difference acting to increase degradation of the mutant mtDNA species. We first estimate a value for ∊_8_ consistent with the heteroplasmy changes involved. Using the transformation

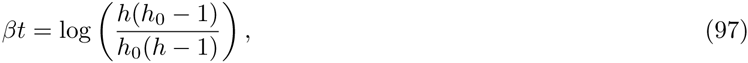

from Eqn. 66 above, where *h*_0_ is initial heteroplasmy and *h* is heteroplasmy at time *t*, we obtain an estimate 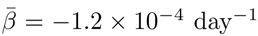. We thus set ∊_8_ = 1.2 × 10^−4^ day^−1^, to produce the required selective difference manifest through mutant degradation.

Solving the ODEs arising from our theoretical approach (Eqns. 34-36) then give values for 〈*w*^2^〉 and 〈*m*^2^〉 over time for a given initial condition. Assuming that each datapoint consists of a sample of 10^3^ cells, we divide these values by 10^3^ to obtain an estimated distribution for each later *w*, *m* pair, given the paired initial *w*, *m* state. We combine these distrbutions to build an overall distribution over later results, and use the Kolmogorov-Smirnov test to test the alternative hypothesis that the later results were incompatible with draws from this distribution. The results were *p* = 0.054 for wildtype mtDNA copy number and *p* = 0.861 for mutant mtDNA copy number. As highlighted in the text, the absence of a *p* < 0.05 result cannot be interpreted as support for the null hypothesis, but this analysis suggests that the available data is not incompatible with the predictions of our model.

## References

[1] G. Capps, D. Samuels, and P. Chinnery. A model of the nuclear control of mitochondrial DNA replication. J. Theor. Biol., 221: 565–583, 2003.

[2] J. St John. The control of mtDNA replication during differentiation and development. BBA, 1840: 1345–1354, 2014.

[3] D. Wallace and D. Chalkia. Mitochondrial DNA genetics and the heteroplasmy conundrum in evolution and disease. CSH Per-spect. Biol., 5: a021220–1 — a021220–49, 2013.

[4] S. Bacman, S. Williams, M. Pinto, S. Peralta, and C. Moraes. Specific elimination of mutant mitochondrial genomes in patient-derived cells by mitoTALENs. Nature Medicine, 19: 1111–1113, 2013.

[5] P. Gammage, J. Rorbach, A. Vincent, E. Rebar, and M. Minczuk. Mitochondrially targeted ZFNs for selective degradation of pathogenic mitochondrial genomes bearing large-scale deletions or point mutations. EMBO Mol. Medicine, 6: 458–466, 2014.

[6] J. Burgstaller, I. Johnston, and J. Poulton. Mitochondrial DNA disease and developmental implications for reproductive strategies. Mol. Hum. Reprod., 21: 11–22, 2015.

[7] R. Rossignol, B. Faustin, C. Rocher, M. Malgat, J. Mazat, and T. Letellier. Mitochondrial threshold effects. Biochem. J, 370: 751–762, 2003.

[8] O. Baris, S. Ederer, J. Neuhaus, J. von Kleist-Retzow, C. Wunderlich, M. Pal, F. Wunderlich, V. Peeva, G. Zsurka, and W. Kunz. Mosaic Deficiency in Mitochondrial Oxidative Metabolism Promotes Cardiac Arrhythmia during Aging. Cell Metabolism, 21: 667–677, 2015.

[9] A. Raj and A. van Oudenaarden. Nature, nurture, or chance: stochastic gene expression and its consequences. Cell, 135: 216–226, 2008.

[10] M. Kærn, T. Elston, W. Blake, and J. Collins. Stochasticity in gene expression: from theories to phenotypes. Nature Rev. Genet., 6: 451–464, 2005.

[11] J. Paulsson. Summing up the noise in gene networks. Nature, 427: 415–418, 2004.

[12] J. Lygeros, K. Koutroumpas, S. Dimopoulos, I. Legouras, P. Kouretas, C. Heichinger, P. Nurse, and Z. Lygerou. Stochastic hybrid modeling of DNA replication across a complete genome. Proc. Natl. Acad. Sci. USA, 105: 12295–12300, 2008.

[13] G. Kopsidas, S. Kovalenko, D. Heffernan, N. Yarovaya, L. Kramarova, D. Stojanovski, J. Borg, M. Islam, A. Caragounis, and A. Linnane. Tissue mitochondrial DNA changes: a stochastic system. Ann. New York Acad. Sci., 908: 226–243, 2000.

[14] I. Johnston, J. Burgstaller, V. Havlicek, T. Kolbe, T. Rulicke, G. Brem, J. Poulton, and N. Jones. Stochastic modelling, Bayesian inference, and new in vivo measurements elucidate the debated mtDNA bottleneck mechanism. eLife, 4: e07464–1 — e07464–44, 2015.

[15] I. Johnston, B. Gaal, R. das Neves, T. Enver, F. Iborra, and N. Jones. Mitochondrial variability as a source of extrinsic cellular noise. PLoS Comput. Biol., 8: e1002416–1 — e1002416–14, 2012.

[16] R. Jajoo, Y. Jung, D. Huh, M. Viana, S. Rafelski, M. Springer, and J. Paulsson. Accurate concentration control of mitochondria and nucleoids. Science, 351: 169–172, 2016.

[17] M. Figge, A. Reichert, M. Meyer-Hermann, and H. Osiewacz. Deceleration of fusion-fission cycles improves mitochondrial quality control during aging. PLoS Comput. Biol., 8: e1002576—1 — e1002576–18, 2012.

[18] S. Poovathingal, J. Gruber, B. Halliwell, and R. Gunawan. Stochastic drift in mitochondrial DNA point mutations: a novel perspective ex silico. PLoS Comput. Biol., 5: e1000572–1 — e1000572–12, 2009.

[19] P. Chinnery and D. Samuels. Relaxed replication of mtDNA: a model with implications for the expression of disease. Am. J. Hum. Genet., 64: 1158—1165, 1999.

[20] I. Johnston and N. Jones. Closed-form stochastic solutions for non-equilibrium dynamics and inheritance of cellular components over many cell divisions. Proc. Roy. Soc. A., 471, 2015.

[21] S. Wright. Statistical genetics and evolution. Bull. Am. Math. Soc., 48: 223–246, 1942.

[22] P. Wonnapinij, P. Chinnery, and D. Samuels. The distribution of mitochondrial DNA heteroplasmy due to random genetic drift. Am. J. Hum. Genet., 83: 582—593, 2008.

[23] J. Burgstaller, I. Johnston, N. Jones, J. Albrechtova, T. Kolbe, C. Vogl, A. Futschik, C. Mayrhofer, D. Klein, S. Sabitzer, M. Blattner, C. Gully, J. Poulton, T. Rulicke, J. Pialek, R. Stein-born, and G. Brem. mtDNA segregation in heteroplasmic tissues is common in vivo and modulated by haplotype differences and developmental stage. Cell Reports, 7: 2031–2041, 2014.

[24] N. Van Kampen. Stochastic processes in physics and chemistry. Elsevier, 1992.

[25] J. Elf and M. Ehrenberg. Fast evaluation of fluctuations in biochemical networks with the linear noise approximation. Genome Res., 13: 2475—2484, 2003.

[26] M. Assaf and B. Meerson. Extinction of metastable stochastic populations. Phys. Rev. E, 81: 021116–1 — 021116–18, 2010.

[27] M. Solignac, J. Genermont, M. Monnerot, and JC. Mounolou. Drosophila mitochondrial genetics: evolution of heteroplasmy through germ line cell divisions. Genetics, 117(4): 687–696, 1987.

[28] T. Wai, D. Teoli, and E. Shoubridge. The mitochondrial DNA genetic bottleneck results from replication of a subpopulation of genomes. Nature Genet., 40: 1484—1488, 2008.

[29] P. Wonnapinij, P. Chinnery, and D. Samuels. Previous estimates of mitochondrial DNA mutation level variance did not account for sampling error: comparing the mtDNA genetic bottleneck in mice and humans. Am. J. Hum. Genet., 86: 540—550, 2010.

[30] M. Kimura. Solution of a process of random genetic drift with a continuous model. Proc. Natl. Acad. Sci. USA, 41: 144–150, 1955.

[31] C. Birky Jr. The inheritance of genes in mitochondria and chloro-plasts: laws, mechanisms, and models. Annu. Rev. Genet., 35: 125–148, 2001.

[32] J. Jenuth, A. Peterson, K. Fu, and E. Shoubridge. Random genetic drift in the female germline explains the rapid segregation of mammalian mitochondrial DNA. Nature Genet., 14: 146–151, 1996.

[33] L. Cree, D. Samuels, S. de Sousa Lopes, H. Rajasimha, P. Won-napinij, J. Mann, H. Dahl, and P. Chinnery. A reduction of mitochondrial DNA molecules during embryogenesis explains the rapid segregation of genotypes. Nature Genet., 40: 249–254, 2008.

[34] L. Cao, H. Shitara, T. Horii, Y. Nagao, H. Imai, K. Abe, T. Hara, J. Hayashi, and H. Yonekawa. The mitochondrial bottleneck occurs without reduction of mtDNA content in female mouse germ cells. Nature Genet., 39: 386–390, 2007.

[35] S. Monnot, N. Gigarel, D. Samuels, P. Burlet, L. Hesters, N. Fry-dman, R. Frydman, V. Kerbrat, B. Funalot, J. Martinovic, A. Benachi, J. Feingold, A. Munnich, and J-P. Bonnefont. Segregation of mtDNA throughout human embryofetal development: m. 3243A? G as a model system. Human Mutation, 32: 116–125, 2011.

[36] K. Lawson and W. Hage. Clonal analysis of the origin of primordial germ cells in the mouse. Germline Dev., 165: 68–84, 1994.

[37] A. Diot, A. Hinks-Roberts, T. Lodge, C. Liao, E. Dombi, K. Morten, S. Brady, C. Fratter, J. Carver, R. Muir, R. Davis, C. Green, I. Johnston, D. Hilton-Jones, C. Sue, H. Mortiboys, and J. Poulton. A novel quantitative assay of mitophagy: Combining high content fluorescence microscopy and mitochondrial DNA load to quantify mitophagy and identify novel pharmacological tools against pathogenic heteroplasmic mtDNA. Pharm. Res., 100: 24–35, 2015.

[38] A. Pyle, R. Taylor, S. Durham, M. Deschauer, A. Schaefer, D. Samuels, and P. Chinnery. Depletion of mitochondrial DNA in leucocytes harbouring the 3243A G mtDNA mutation. J. Med. Genet., 44: 69–74, 2007.

[39] I. Johnston, B. Rickett, and N. Jones. Explicit tracking of uncertainty increases the power of quantitative rule-of-thumb reasoning in cell biology. Biophys. J., 107(11): 2612–2617, 2014.

[40] I. Johnston. Efficient parametric inference for stochastic biological systems with measured variability. Stat. Appl. Genet. Mol. Biol., 13: 379–390, 2014.

[41] K. Åström. Introduction to stochastic control theory. Dover, 2012.

[42] V. Borkar. Controlled diffusion processes. Probability Surveys, 2: 213–244, 2005.

[43] H. Markowitz. Portfolio selection. J. Finance, 7: 77–91, 1952.

[44] D. Gillespie. Exact stochastic simulation of coupled chemical reactions. J. Phys. Chem., 81: 2340–2361, 1977.

[45] J. Jenuth, A. Peterson, and E. Shoubridge. Tissue-specific selection for different mtDNA genotypes in heteroplasmic mice. Nature Genet., 16: 93–95, 1997.

[46] R. Jokinen, P. Marttinen, H. Sandell, T. Manninen, H. Teeren-hovi, T. Wai, D. Teoli, J. Loredo-Osti, E. Shoubridge, and B. Battersby. Gimap3 regulates tissue-specific mitochondrial DNA segregation. PLoS Genet., 6: e1001161–1 – e1001161–9, 2010.

[47] T. Toni, D. Welch, N. Strelkowa, A. Ipsen, and M. Stumpf. Approximate Bayesian computation scheme for parameter inference and model selection in dynamical systems. J. Roy. Soc. Interf., 6: 187–202, 2009.

[48] http://www.bio-rad.com/webroot/web/pdf/lsr/literature/4106212b.pdf.

